# Where Honey Bee Vitellogenin may Bind Zn^2+^-Ions

**DOI:** 10.1101/2022.01.28.478200

**Authors:** Vilde Leipart, Øyvind Enger, Diana Cornelia Turcu, Olena Dobrovolska, Finn Drabløs, Øyvind Halskau, Gro V. Amdam

**Affiliations:** Faculty of Environmental Sciences and Natural Resource Management, Norwegian University of Life Sciences, Aas, Norway; Department of Biological Sciences, University of Bergen, Bergen, Norway; Helse Bergen, Bergen, Norway; Department of Clinical and Molecular Medicine, Faculty of Medicine and Health Sciences, NTNU – Norwegian University of Science and Technology, Trondheim, Norway; School of Life Sciences, Arizona State University, Tempe, AZ, United States

**Keywords:** Honey bees, vitellogenin, zinc-binding, insect immunity, protein structure analysis

## Abstract

The protein Vitellogenin (Vg) plays a central role in lipid transportation in most egg-laying animals. High Vg levels correlate with stress resistance and lifespan potential in honey bees (*Apis mellifera*). Vg is the primary circulating zinc-carrying protein in honey bees. Zinc is an essential metal ion in numerous biological processes, including the function and structure of many proteins. Measurements of Zn^2+^ suggest a variable number of ions per Vg molecule in different animal species, but the molecular implications of zinc-binding by this protein are not well understood. We used inductively coupled plasma mass spectrometry (ICP-MS) to determine that, on average, each honey bee Vg molecule binds 3 Zn^2+^-ions. Our full-length protein structure and sequence analysis revealed seven potential zinc-binding sites. These are located in the β-barrel and α-helical subdomains of the N-terminal domain, the lipid binding site, and the cysteine-rich C-terminal region of unknown function. Interestingly, two potential zinc-binding sites in the β-barrel can support a proposed role for this structure in DNA-binding. Overall, our findings illustrate the capacity of honey bee Vg to bind zinc at several functional regions, indicating that Zn^2+^-ions are important for many of the activities of this protein. In addition to being potentially relevant for other egg-laying species, these insights provide a platform for studies of metal ions in bee health, which is of global interest due to recent declines in pollinator numbers.

## Introduction

Zinc is necessary for living organisms to function properly (Sloup et al., 2017). The element is involved in basic life processes such as cell division and gene expression and it is an essential nutrient for growth and development (Falchuk, 1998, Baltaci and Yuce, 2018). Zinc is necessary for catalytic, structural, and regulatory functions for thousands of proteins (Andreini et al., 2006). Improving the understanding of zinc-carrying proteins is thus likely to reveal new information about many physiological processes across taxa.

An important zinc-carrying protein in egg-laying animals is the multi-domain glycolipophosphoprotein Vitellogenin (Vg) (Montorzi et al., 1994, Falchuk, 1998, Matozzo et al., 2008, Gupta et al., 2021). Vg provides nutrients to developing embryos by delivering lipids, amino acids, and zinc (Pan et al., 1969). In some species, Vg is expressed in juveniles, as well as in males and in females that do not reproduce, hinting at roles beyond yolk formation (Sappington and S. Raikhel, 1998). Such roles have been most abundantly studied in honey bees (*Apis mellifera*), where Vg is recognized as a multi-functional protein impacting the behavior and health of workers (functionally sterile females). RNA-interference mediated gene knockdown reveals that honey bee Vg affects worker behavioral ontogeny, foraging choice, capacity to provide larval care, stress resistance, and longevity (Amdam et al., 2004, Guidugli et al., 2005). At least some of the physiological impact of honey bee Vg can be zinc-related, as low zinc levels may reduce the cell-based immune capacity of the workers (Amdam et al., 2005). However, apart from the finding that circulating zinc levels correlate strongly with the hemolymph level of Vg in honey bees, the molecular relationship between this ion and protein is largely unknown.

The capacity for zinc-binding is established for Vg proteins, but the number of ions per Vg molecule varies among species. For example, measurements of bound Zn^2+^-ions in the hemolymph of shore crab (*Carcinus maenas*) (Martin and Rainbow, 1998) are higher than those of the American clawed frog (*Xenopus laevis*) (Montorzi et al., 1994) and domestic fowl (*Gallus gallus*) (Mitchell and Carlisle, 1991). The same studies also indicate that the number of Zn^2+^-ions carried by Vg can vary with individual reproductive state and age. Such variation is likely biologically important, but to date is not well understood for Vg proteins.

A prerequisite for understanding zinc-related molecular mechanisms is finding the location and structural context of Zn^2+^-binding sites within the protein of interest (Ataie et al., 2008, Daniel and Farrell, 2014). The coordinating environment for Zn^2+^-ions in proteins is well characterized (Dudev and Lim, 2003, Pace and Weerapana, 2014), and binding sites are usually sorted into two structurally distinct categories based on whether the Zn^2+^ has a catalytic or structural role. A catalytic binding site is often partially exposed to the solvent, and the Zn^2+^-ion coordinates most often with histidine (H), cysteine (C), aspartate (D), glutamate (E), serine (S) residues, and/or water molecules (Jernigan et al., 1994, Ataie et al., 2008). A structural binding site is usually buried in the protein, surrounded by an intricate network of hydrogen bonds (Dudev and Lim, 2003), and the Zn^2+^-coordinating residues are typically multiple H/C residues only. For example, the well-known transcription factor motif zinc fingers (C4 or C2H2) (Pace and Weerapana, 2014) represents a structural binding site.

The prevalence of coordinating residues differs between catalytic and structural binding sites. For catalytic sites, 4, 5, and 6 residues coordinate in 48, 44, and 6 % of cases, respectively. Correspondingly, the ratio is 79, 6, and 12 % for structural sites, respectively (Dudev and Lim, 2003, Ataie et al., 2008). These numbers imply that catalytic sites largely coordinate Zn^2+^ with 4 or 5 residues, while structural sites most commonly coordinate with 4 residues. In addition to these two categories, Zn^2+^-binding is identified in regulatory Zn^2+^-Cys complexes, called redox switches (Pace and Weerapana, 2014). Two common characteristics for all coordinating sites are: strong interaction between the residues and the Zn^2+^-ion, and high hydrophobic contrast in the binding site (Dudev and Lim, 2003, Pace and Weerapana, 2014).

The zinc coordinating environment for honey bee Vg is not described, but speculations have been presented regarding lamprey (*Ichthyomyzon unicuspis*) and zebrafish (*Danio rerio*) Vg. Anderson *et al*. (1998) published the only experimentally solved protein structure of lamprey Vg (PDB-ID: 1LSH), which lacks Zn^2+^-ions due to use of 1 mM EDTA during crystallization. In the absence of zinc, Anderson *et al*. (1998) proposed two potential binding sites (H312/H322 and H868/H887) based on the residues’ locations in the crystal structure and the sequence conservation. In comparison, Sullivan *et al*. (2018) suggest the phosphorylated serine-rich phosvitin domain is associated with Zn^2+^-ions in zebrafish (Sullivan and Yilmaz, 2018). This domain is missing in some Vg proteins, including those of insects (Tufail and Takeda, 2008). For example, Honey bee Vg consists of an N-terminal domain, a lipid cavity, and a C-terminal region (Havukainen et al., 2011, Havukainen et al., 2012, Leipart et al., 2022). The N-terminal domain comprises the β-barrel subdomain followed by a flexible polyserine linker and the α-helical subdomain. The lipid cavity is built up by a domain of unknown function (DUF1943), a β-sheet, and a von Willebrand factor (vWF) domain. Honey bee Vg can be cleaved at the polyserine linker in the N-terminal domain. This cleavage creates a small fragment (40 kDa, the β-barrel subdomain) and a larger fragment (150 kDa, the α-helical subdomain, the lipid binding site, and the C-terminal region) (Havukainen et al., 2012). It is interesting to note that the smaller fragment of honey bee Vg may translocate into cell nuclei, bind DNA (potentially with co-factors), and influence gene expression (Salmela et al., 2021).

Honey bees provide a practical and useful research system as they are globally available as commercial pollinators and primary producers of honey, pollen, and wax. We recently published the first full-length Vg structure prediction for honey bees (Leipart et al., 2022) using experimental available data, computational modeling and AlphaFold. The structural templates have not resolved the position of zinc, and AlphaFold does not predict the position of non-protein components (Jumper et al., 2021). We use the model here to provide insight into knowledge of where Vg binds zinc. First, we performed an element analysis of Vg protein obtained from worker bee hemolymph and found that, on average, it binds 3 Zn^2+^-ions. Using structural data in combination with sequence data, we then conducted an in-depth analysis to predict the location(s) of potential zinc-binding sites. We identified areas in the β-barrel subdomain, α-helical subdomain, lipid binding site, and C-terminal region. We propose that a zinc-binding site in the β-barrel subdomain plays a role in DNA binding. In an attempt to characterize the zinc-binding site(s) of this subdomain, we expressed the β-barrel using bacterial recombinant expression systems in cultural medium with various compositions of Zn^2+^ and/or Co^2+^ but this approach did not provide a clear answer. However, taken together, our results provide the first detailed insights into where honey bee Vg can bind Zn^2+^.

## Results

### Identification of zinc in honey bee Vg using inductively coupled plasma mass spectrometry

We performed inductively coupled plasma mass spectrometry (ICP-MS) on Vg from worker bee hemolymph to confirm Vg as a zinc carrier and quantify the number of Zn^2+^-ions per Vg molecule. We detected significant amounts of Zn^2+^ in Vg samples relative to the (zinc negative) controls (Kruskal-Wallis analysis: chi-squared = 7.81, df = 1, p-value = 0.00519, Figure 1A). Based on sample concentrations of Vg and Zn^2+^, and using the theoretical molecular weight of Vg and Zn^2+^ (201147.7 g/mol and 65.30 g/mol, respectively), we calculated the molecular Zn:Vg ratio for each sample (Figure 1B). This analysis gave a range of 2.57–3.89 mol of Zn^2+^-ions per Vg molecule, with an average ratio of 3 Zn^2+^-ions per full-length Vg protein.

**Figure 1.**
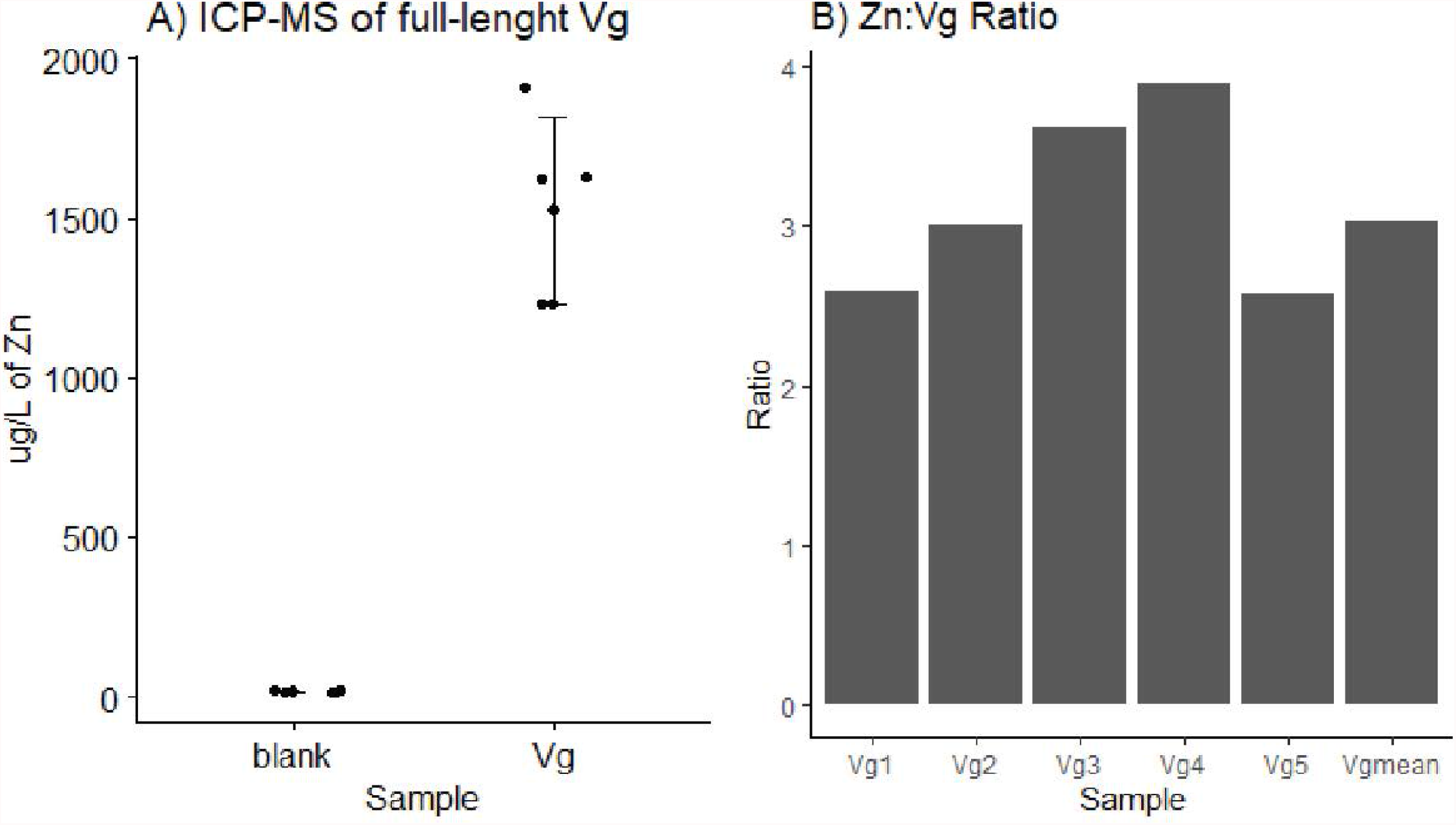
ICP-MS. **A)** The concentration measured with ICP-MS for the 5 blank and full-length Vg samples is plotted. The mean and the standard deviation of the mean are indicated for each group (lines). **B)** The calculated molecular Zn:Vg ratio for each sample of Vg is provided, and the calculated mean value for all samples is included as a separate bar.

### Identification of potential zinc coordinating residues in honey bee Vg

In this section, we analyze honey bee Vg to identify zinc-binding sites, referred to as clusters. The results are summarized in Table 1 and illustrated in Figure 2A. To achieve this, we took a comprehensive approach: using online bioinformatic tools developed for identification of zinc motifs in amino acid sequences, assessing suggested structural sites from studies on lamprey (Anderson et al., 1998) and zebrafish (Sullivan and Yilmaz, 2018), and finally, fully analyzing our recently published protein structure (Leipart et al., 2022). We used a multiple sequence alignment (MSA) of Vg sequences with a broad phylogenetic range to evaluate the findings.

**Table 1:** Identified zinc clusters.

| Cluster | Vg domain | Residues |
| --- | --- | --- |
| βb.1 | β-barrel subdomain | H20, H113, D143, E147, H265 |
| βb.2 | β-barrel subdomain | E171, D172, S173, C178, E179, C222, D223 |
| αh.1 | α-helical subdomain | H229, H577, H587, H593, H602 |
| αh.2 | α-helical subdomain | H697, E698, C701, S775, S800, D802 |
| Duf.1 | DUF1943 | E987, H988, H990, H1045, D1046 |
| Duf.2 | DUF1943 | H445, E449, D996, H1000, H1035, S1037 |
| Ct | C-terminal | C1687, C1711, C1715, C1768 |

The clusters are named (column 1) based on their location in honey bee Vg (column 2). The residues included in each cluster are listed in column 3.

**Figure 2.**
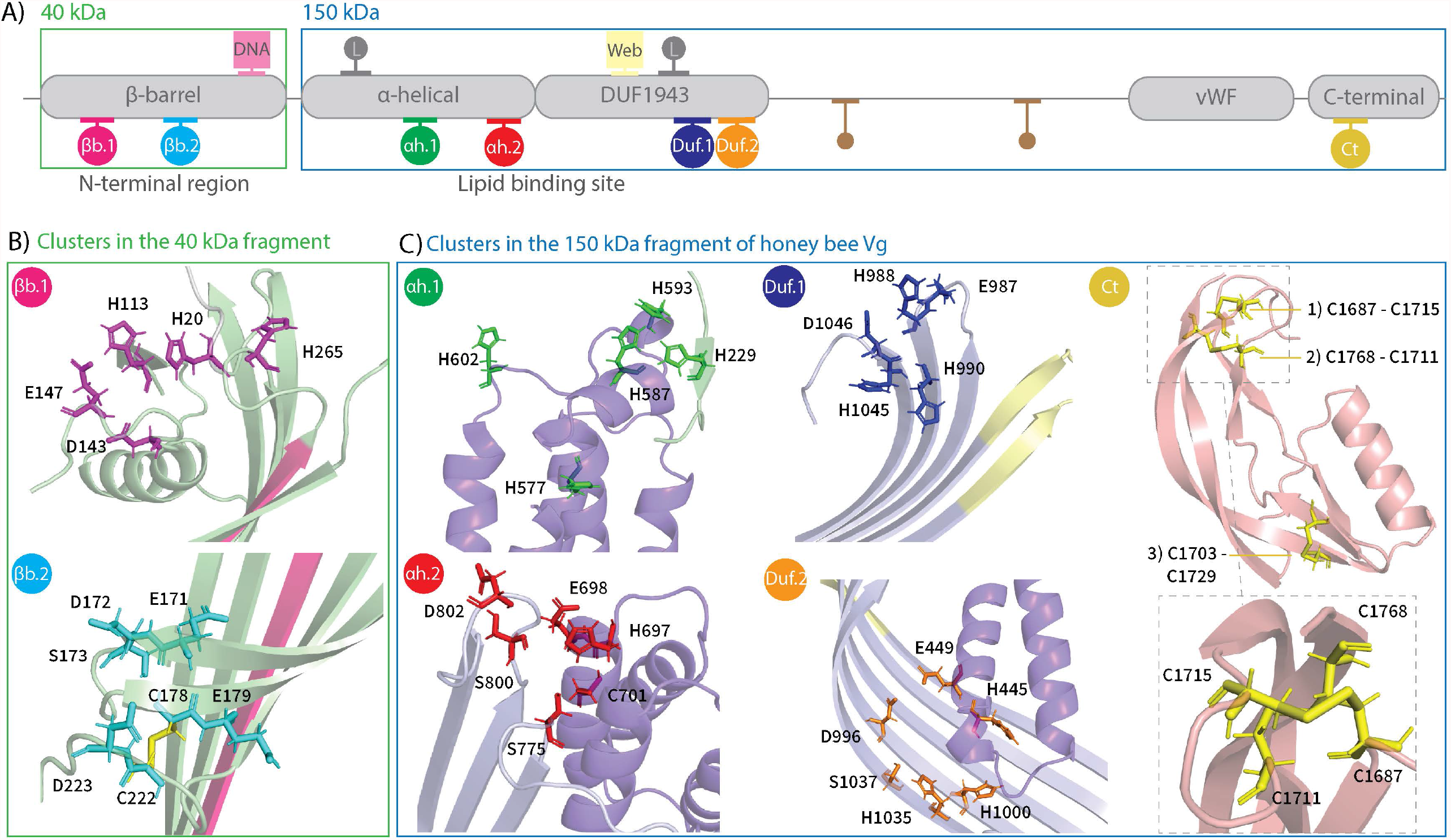
Identified clusters: **A)** 2D representation of the honey bee Vg with conserved domain, subdomains, and regions (gray boxes). The amino acid sequence was divided into two regions based on the naturally cleaved fragments, the α-helical subdomain, the lipid binding site, including vWF and C-terminal region (blue, 150 kDa), and the β-barrel subdomain (green, 40 kDa). The identified clusters are marked below the gray boxes (αh.1: green, αh.2: red, Duf.1: blue, Duf.2: orange, Ct: yellow, βb.1: pink and βb.2: cyan). The two zinc-binding locations identified by studies of lamprey are marked above the gray boxes with gray dots (L). The DNA binding motif (pink box, DNA) and the zinc-binding motif identified by MotifScan (yellow box, web) are also marked above the gray boxes. Finally, the two conserved disulfide bridges not included in a cluster are included here (smaller brown dots). **B)** Identified clusters in the 40 kDa fragment of honey bee Vg. The residues of cluster βb.1 (magenta) and βb.2 (cyan) are in the β-barrel subdomain (green) and close to the DNA binding motif (pink β-sheet). The disulfide bridge in cluster βb.2 is shown as a yellow stick. **C)** Identified clusters in the 150 kDa fragment of honey bee Vg. <u>Cluster αh.1:</u> The insect-specific loop region in the α-helical subdomain (purple) and a short β-strand from the β-barrel subdomain (green) contains the residues of cluster αh.1 (green sticks). <u>Cluster αh.2:</u> The α-helical subdomain (purple) and the β-sheet region in the DUF1943 (light purple) are shown as a cartoon. The identified cluster-residues are shown as red sticks. <u>Cluster Duf.1:</u> Two loops from the DUF1943 (light purple) contain the identified residues, shown as blue sticks. The zinc-binding motif identified by MotifScan (yellow) is located in the neighboring β-strands in the DUF1943. <u>Cluster Duf.2:</u> Cluster Duf.2 (orange sticks) is found further down on the same β-sheet in DUF1943 (light purple). Two of the residues are located in the α-helical subdomain (purple). <u>Cluster Ct:</u> Three disulfide bridges (yellow sticks) are found in the C-terminal (light pink). The dotted box zoom on two of the disulfide bridges shows them from another direction. The two disulfide bridges can create a tetrahedral geometry and are identified as cluster Ct.

#### Assessing potential zinc coordinating residues using online tools

The searches using online motif search algorithms (see Experimental procedures) resulted in only one hit. MotifScan (Pagni et al., 2007) identified a Zn^2+^-binding motif at residue 926 to 936. The short region includes several conserved residues (F928, P929, G933, L934, P935, and F936, Figure S1). Regarding the MSA (Figure S1), the zinc-binding residue identified in the motif, H926, is only present in the *Apis mellifera* Vg sequence. The motif is located in a β-strand-turn-β-strand fold in the DUF1943, which extends into the β-barrel subdomain, close to cluster Duf.1 and βb.2, and the DNA binding motif (see Figure S2 for a structural overview of this region).

#### Assessing potential zinc coordinating residues from suggested zinc-binding sites

According to the MSA, the two sites proposed in lamprey Vg (H312/H322 and H868/H887) align to E427/V444 and N975/G994 in honey bee Vg, respectively (Figure S1B). Of the four residues in honey bee Vg, only the E residue is known to coordinate with Zn^2+^. The low conservation of E427 and lack of other typically Zn^2+^-coordinating residues (H/C/D/S) at both sites suggest that these locations do not take part in zinc-binding in the honey bee.

Sullivan and Yilmaz (2018) suggest that the phosvitin domain in zebrafish Vg has a Zn^2+^-coordinating role. Honey bee Vg does not have this domain, but the polyserine linker in the N-terminal domain is a similar, serine-rich region. We did not identify any C, H, D, or E residues here that could support zinc-coordination, but the linker contains 14 S residues. One or two S residues can participate in zinc-coordination. However, this location is an unlikely candidate as it would be unprecedented to find Zn^2+^-coordination by serines in a disordered (loop) region (Baglivo et al., 2009).

#### Assessing potential zinc coordinating residues based on protein structure modeling

##### Potential Zn^2+^ sites in the β-barrel subdomain of the N-terminal region

The N-terminal β-barrel subdomain contains a total of seven H residues (H20, H47, H113, H193, H210, H229, and H265) and two C residues (C178 and C222) (Figure 2B). Among these, H229 is not folded in the β-barrel subdomain (see cluster αh.1 in Figure 2C). The C residues are conserved in most of the Vg sequences included in the MSA (Figure S1A and Figure S3A). H20 and H113 are conserved in the Vg sequences in the MSA from insect species, while the remaining H residues are less conserved. The first cluster identified in the β-barrel subdomain, βb.1, contains H20 and H113 located in separate loop regions (Figure 2B). H265 is situated at the beginning of a β-strand close to βb.1. In addition, we found two conserved residues (D143 and E147) in a loop region and included them in cluster βb.1 (Table 1).

The second cluster in the β-barrel subdomain, βb.2, contains C178 and C222. The C residues form a disulfide bridge. We identified five conserved residues (E171, D172, S173, E179, and D223) close to the disulfide bridge and included them in cluster βb.2. All residues are in two neighboring β-strands, apart from C222 and D223 at the end of a loop region (Table 1, Figure 2B).

A short SRSSTSR sequence (residues 250 to 256) in the β-barrel subdomain is proposed to bind DNA (Salmela et al., 2021). The residues are located at a β-strand close to cluster βb.1 and βb.2 (Figure S2 and Figure S3A).

##### Potential Zn^2+^ sites in α-helical subdomain and lipid binding site

We identified four clusters of conserved H/C residues (Table 1), two in the α-helical subdomain (αh.1 and αh.2) and two in the DUF1943 (Duf.1 and Duf.2) (Figure S2C and Figure S1B). The first cluster in the α-helical subdomain, αh.1, contains two highly conserved H residues (H587 and H593) and two less conserved H residues (H577 and H602). All H residues are in a well-conserved insect-specific loop in the α-helical subdomain (Figure S3B). The cluster also includes the well-conserved H229. The residue is part of a loop extending from the β-barrel subdomain, which positions H229 close to H587 and H593. The second cluster in the α-helical subdomain, αh.2, is located at the beginning of the well-conserved 15^th^ α-helix in the subdomain (Figure S3B) containing H697, C701, and E698 (Figure 2C). The cavity-facing side of the α-helical subdomain is close to a β-sheet region of DUF1943. More specifically, the α-helices are near two loops connecting the β-strands on DUF1943. We identified three well-conserved residues in those loops (S775, S800, and D802) and included them in cluster αh.1.

The first cluster in the DUF1943, Duf.1, contains three conserved H residues (H988, H990, and H1045). They are packed together in two loop regions at the end of two adjacent β-strands (Figure 2C). Two conserved residues (E987 and D1046) were identified in the same loops and included in Duf.1. The second cluster in DUF194, Duf.2, contains two conserved H residues (H1000 and H1035) positioned on the same β-sheet as Duf.1 (Figure S1B and Figure S3C). We also found two conserved residues (D996 and S1037) in the β-sheet region and two residues in the α-helical subdomain (H445 and E449 in the second α-helix of the subdomain) and included those in Duf.2.

##### Potential Zn^2+^ sites in lipid binding site β-sheet and the C-terminal region

Following the DUF1943, we identified four conserved C residues (C1242, C1279, C1310, and C1324) in a β-sheet in the lipid binding site (Figure S1C). C1242 and C1279 are highly conserved and create a disulfide bridge in the β-sheet region. C1310 and C1324 also create a disulfide bridge connecting two α-helices. Neither of the disulfide bridges has any conserved D, E, or S residues in proximity, so the bridges are not identified as potential zinc clusters here. After this β-sheet in the lipid binding site, the following domain is the vWF domain, which does not coordinate Zn^2+^ (Leipart et al., 2022).

The final region in honey bee Vg is the C-terminal (residue 1635 to 1770). This region contains seven C and two H residues that are conserved in most Vg sequences in the MSA (Figure S1D and Figure S3D). Six C residues create three separate disulfide bridges (Figure 2C). Two of these bridges cross each other (C1687, C1711, C1715, and C1768) and were identified as cluster Ct (Table 1). The remaining disulfide bridge and conserved C and two H residues were not nearby and therefore not considered a potential zinc-binding site.

#### Assessing functional roles of specific zinc coordinating residues

##### Motif comparison in the β-barrel subdomain

The functional roles of zinc-binding sites are not well defined for any Vg. However, a possible exception is made by the short SRSSTSR sequence close to clusters βb.1 and βb.2 in the β-barrel subdomain that may bind DNA in honey bees (Figure 2B and S2). A recent study presents a proposed DNA-binding motif that the subdomain recognizes (see Figure 5 in Salmela *et al*. 2021). Motif A is similar to the transcription factor CTCF motifs in *Drosophila melanogaster* (see Figure S5A and S5B for motif comparison). CTCF contains eleven C2H2 zinc finger factor motifs (Maksimenko et al., 2021) that bind one Zn^2+^-ion each, shown to stabilize the structural fold in CTCF required to bind DNA (Maksimenko et al., 2021). We aligned the C2H2 motifs to the Vg sequences in our MSA (Figure S5C) and found five predicted zinc-binding residues in the β-barrel subdomain aligned to C and H residues in the C2H2 motifs. D143 and E147 from βb.1 align to H and D residues in the C2H2 motifs, and C178 and C222 from βb.2 align to C residues in the C2H2 motifs. In addition, H229 from αh.1 aligns to H residues in the C2H2 motifs. This configuration of zinc-binding residues in honey bee Vg supports a functional role of the β-barrel in DNA binding and, more specifically, suggests that Vg needs at least one Zn^2+^-ion to stabilize the binding to DNA.

##### Attempting detection of zinc in the β-barrel subdomain

To begin validating the role of zinc in the honey bee Vg β-barrel subdomain, we initially attempted to obtain a sample through dephosphorylation and proteolysis of native Vg at the polyserine linker. However, this was unsuccessful. A weak 40kDa band was obtained under certain conditions, but we could not validate its identity (see Figure S6 for SDS-PAGE gel). Therefore, the subdomain was expressed in *E. coli* with a solubility tag (SUMO) by Genscript Biotech, and ICP-MS was repeated. The Zn^2+^ concentration for the tagged β-barrel subdomain samples was significantly higher than the negative controls of buffer samples (Mann-Whitney U test: w = 0, p-value = 0.0106) (see Figure S7 for ICP-MS results), but not significantly different from samples of the free SUMO-tag, including all samples (Mann-Whitney U test: w = 4, p-value = 0.0937). Excluding one outlier in the SUMO-tag sample set, however, yielded significant results (Mann-Whitney U test: w = 0, p-value = 0.0195). This outcome seems encouraging, but the tagged β-barrel subdomain was exposed to Zn^2+^ during expression in culture, while the free SUMO-tag was synthetically produced without similar opportunity for zinc-binding. We attempted to develop systems to control for this confounding factor (see supplementary.docx and Figure S7 and S8), but without sufficient success.

## Discussion

Our study validates that honey bee Vg binds Zn^2+^, as suggested previously (Amdam et al., 2004). The Zn:Vg ratio calculated by us, 3:1, is higher than the 1 or possible 2 zinc ions reported for each monomer of lamprey Vg (Anderson et al., 1998), which was a calculation based on measurements from the American clawed frog Vg (Montorzi et al., 1994, Auld et al., 1996). After the initial validation, we performed a sequence and structural analysis to understand the structural basis and possible functional outcomes of the zinc-binding capacity of honey bee Vg. We identified seven zinc clusters located at different subdomains and domains.

We found two clusters in the β-barrel subdomain. The β-barrel subdomain is proposed to function as a transcription factor (Salmela et al., 2021). A classical feature in the DNA binding domain of transcription factor is the coordination of Zn^2+^, called zinc finger domains (Cassandri et al., 2017). The Zn^2+^-binding assists the protein in folding, creating the structural form that can recognize DNA (Chang et al., 2010). We show a similarity between a known zinc finger protein and honey bee Vg in the DNA binding motif. The CTCF transcription factor requires coordination of Zn^2+^ at the C2H2 motifs to adopt the correct fold to bind DNA (Maksimenko et al., 2021). Similar to CTCF, the β-barrel subdomain might also bind zinc, and through this process build a different fold than seen in our model. The MSA (Figure S5C) shows two C residues in cluster βb.2 and H229 in the β-barrel subdomain, aligned to C and H zinc-binding residues in CTCF. The three residues are at distant positions in the subdomain (Figure S2). The β-barrel subdomain is presumably cleaved from honey bee Vg when translocating to the nucleus (Salmela et al., 2021). Proteolytic cleavage makes H229 available (no longer associated in the αh.1 cluster). The loop region of H229 allows for flexibility and possible association with the two C residues in cluster βb.2, creating a C2H1 site. Cluster βb.1 consists of three H residues in loop regions, which could fold similarly, resulting in a C2H2 zinc site. Taken together, we demonstrated that the β-barrel subdomain can potentially create a C2H2 zinc site if flexibility in the loop regions is allowed for. Zinc-binding-related flexibility is documented for several C2H2 zinc-binding factors, in addition to CTCF, in insects (Jauch et al., 2003, Stubbs L. et al., 2011, Maksimenko et al., 2021). Such flexibility could be interrupted by the bound SUMO-tag in our expression system. In addition, the CTCF transcription factor consists of several C2H2 motifs linked together when binding DNA (Maksimenko et al., 2021, Jauch et al., 2003). Similarly, the β-barrel subdomain might require co-factors or proteins to bind DNA (Salmela et al., 2021).

Cluster αh.1 is in a histidine-rich loop in the α-helical subdomain, a loop identified earlier as flexible and insect-specific in honey bee Vg (Havukainen et al., 2013). A structural zinc-binding site could increase the stability of the α-helical subdomain by structuring the loop and stabilizing the N-terminal domain through interaction with H229 (Figure 2C). However, proteolytic cleavage would disengage H229 from the site. We suggest that the remaining four H located in the loop (Figure 2C) could create a new structural zinc site in such situations, similar to some zinc transporter proteins (Fukada and Kambe, 2011, Tanaka et al., 2013, Zhang et al., 2019). We suggest that a conformational change induced by zinc is feasible for the flexible loop and could stabilize the α-helical subdomain and, in turn, the lipid binding site. Cluster αh.1 is at the surface of honey bee Vg. We propose that the αh.1 cluster could sense the cellular environment more efficiently than the clusters inside the lipid cavity. Zinc transporters with similar histidine-loop coordination sites regulate zinc homeostasis (Fukada and Kambe, 2011), supporting our findings.

Electrostatic and hydrophobic interactions between the α-helices in the α-helical subdomain and two β-sheets in the DUF1943 stabilize the Vg lipid cavity (Babin et al., 1999, Smolenaars et al., 2007, Biterova et al., 2019). We suggest that structural zinc-binding sites in the cavity could support structural stability for lipid uptake and delivery (Smolenaars et al., 2007). Clusters αh.2, Duf.1, and Duf.2 have residues at the α-helical subdomain and the DUF1943. The clusters are in conserved (Figure S3B and S3C) and hydrophobic regions (see Figure S4 for a structural overview of the hydrophobic areas), and Duf.1 and Duf.2 consist of three H residues, typical at a structural zinc site (Dudev and Lim, 2003). Cluster αh.2 has only one H residue but has an additional C residue, a rare arrangement for a structural site, but identified in a ubiquitin-binding protein ((Lim et al., 2019) PDB-ID: 6H3A), indicating that just two H/C residues could coordinate zinc in cluster αh.2. The structural zinc-binding sites in zinc transporters can have an additional D residue (Fukada and Kambe, 2011), which suggests a possibility for the structural coordination event in cluster αh.2 to include a D residue (D802).

We speculated whether an interaction between H926 at the DUF1943 and the two C residues in cluster βb.2 (Figure S2) is possible. However, the residues are too distant (∼30 Å) for interaction (normally ∼2.0–2.4 Å (Dudev and Lim, 2003)) in our model. The generally rigid β-sheets (Perczel et al., 2005) at both positions make a conformational change induced by zinc unlikely. H926 and Duf.1 are at the same β-sheet. However, a similar rigidness would also make interaction unlikely. Therefore, H926 is less likely to coordinate zinc, while cluster αh.2 and Duf.1, Duf.2 are feasible structural zinc-binding sites in the lipid cavity and can coordinate zinc (e.g., during transport). We suggest an optimal solution for honey bee Vg would be to carry lipid molecules and zinc in the same location so it could be released together upon delivery.

The well-conserved disulfide bridge on the C-terminal region presumably contributes to a stable structural fold. Such bridges are usually stable oxidative conditions(Sevier and Kaiser, 2002). Two of the disulfide bridges create an interesting arrangement and suggest the possibility for a ZnC4 coordination site, which would probably maintain the stable fold when Vg is experiencing reducing conditions. ZnC4 is a typical coordination site for redox switches (Ilbert et al., 2006, Pace and Weerapana, 2014). Redox switches can sense oxidative stress, which could generate a response of the protein to change cellular location or release zinc (Ilbert et al., 2006). We propose a similar process that can subside at the Ct cluster in honey bee Vg. This idea has some support in the observation that Vg levels in worker honey bees are positively correlated with oxidative stress resilience (Seehuus et al 2006). It is possible that the zinc released from Vg can explain this phenomenon via binding to protective enzymes or cell membranes (Marreiro et al., 2017, Seehuus et al., 2006).

Regarding the caveats of this study, we assert that *in silico* analysis of the β-barrel subdomain is not fully reflected in our experimental results, despite using several methodologies to approach this problem. Proteolytic cleavage of the SUMO-tag resulted in low yields, and separation during purification of the tag-free subdomain from the SUMO-tag did not work, despite optimization. Changing the solubility-tag to maltose-binding protein (MBP) improved expression yields slightly. However, we faced the same challenges as earlier during purification. Adding two affinity column purification steps followed by a size exclusion chromatography did not successfully separate the SUMO-tag from the subdomain. The tagged subdomain, therefore, became the best option for element analysis. Another drawback of *in silico* prediction is that the orientation for some side chains can be imprecise, even when located in a confident backbone fold (Jumper et al., 2021). The residues in cluster αh.2, H593, H229, and H587 look to have an optimal side chain arrangement to coordinate zinc, creating a small triangle (Figure 2C). However, the side chains might not always be in such an optimal arrangement as seen, for example, in Duf.1. Specifically, the side chain orientation can be inaccurately predicted or could adopt another side chain orientation when zinc is present (Kluska et al., 2018). Due to these caveats, we identified the residues as potential clusters. We also assumed this for cluster βb.1, βb.2, αh.2, and Duf.2 (Figure 2B-C).

Our analysis relied on our recently published full-length protein structure that was predicted using AlphaFold (Leipart et al., 2022). AlphaFold calculates a confidence score that evaluates the predicted structure’s stereochemical integrity (Mariani et al., 2013, Jumper et al., 2021). The honey bee Vg prediction has a confidently folded backbone, with the exception of a few short loops and regions that make up approximately 13% of the Vg residues (see Figure 2 in (Leipart et al., 2022)). The zinc clusters that we identified here are fully embedded in confidently folded regions. However, loop regions are flexible structures (Barozet et al., 2021) and residues located in such flexible regions could potentially change the backbone fold when zinc is present, as seen in zinc regulators and zinc transporter proteins (Liu et al., 2021, Tanaka et al., 2013, De Angelis et al., 2010). We assume this could be true in our model, and this insight was applied in clusters βb.1, αh.1, αh.2, and Duf.1 (Figure 2B-C).

While the localization of one or more zinc-binding sites to the recombinantly expressed β-barrel remains uncertain, the presence of, on average, 3 Zn^2+^ cations per honey bee Vg molecule was determined with confidence using ICP-MS (Figure 1). Our structural analysis illustrates where Vg can bind zinc ions, presumably one at each position. The seven sites might provide unknown flexibilities based on Vg having Zn^2+^-ions bound at different combinations of sites depending on the protein’s situation.

## Experimental procedures

### Collection and purification of honey bee Vg

To obtain Vg, we collected 1-10 µl honey bee hemolymph in a 1:10 dilution in 0.5 M Tris HCl, using BD needles (30G), as described earlier (Aase et al., 2005). The dilution was filtered using an 0.2 µm syringe filter. Vg was purified from honey bee hemolymph with ion-exchange chromatography using a HiTrap Q FF 1mL column. The sample buffer (0.5 M Tris HCl) and elution buffer (0.5 M Tris HCl with 0.45 M NaCl) was prepared with ion-free water and acid-treated (10% HNO_3_ overnight) plastic bottles to eliminate zinc contamination. Then 400–450 µl diluted hemolymph was manually injected and Vg eluted at the conductivity of 15–22 mS/cm. All fractions from the peak were collected, pooled, and up-concentrated using an Amicon® Ultracel 100 kDa membrane centrifuge filter. We verified the fraction purity by running SDS-PAGE, which contained one band only of the correct size (∼180 kDa). The purification protocol was repeated to produce five samples with concentrations between 1.2 and 2.8 µg/µl in 65 µl. Five blank samples were created using the same protocol; here, only sample buffer was injected. The protein concentration, measured with Qubit, confirmed the samples contained no protein. All samples were collected in 1.5 mL Eppendorf tubes pretreated with 10% HNO_3_ for 24 hours and dried at 65 °C.

### Identification of zinc in honey bee Vg using Inductively Coupled Plasma mass spectrometry

ICP-MS was performed to detect metal ions associated with purified Vg. For the ICP-QQQ-MS of full-length Vg extracted from bees, 32 µL of concentrated ultra-pure(up) nitric acid was added to the 65µL samples (purified Vg and blanks as described above). The samples were placed in a heating cabinet at 90°C for 3 hours and subsequently put into an ultrasound bath for 60 seconds to dissolve any remaining particles before analysis. Then the samples were diluted to 325 µL by adding a solution of 1% (V/V) HNO_3_ and 28.5 µg/L Rhodium (Rh, used as an internal standard for Zn), an x5 total volume dilution. This gives a Rh concentration of 20 µg/L in the final sample. For ICP-QQQ-MS analysis on the recombinantly expressed and purified β-barrel, 30 µL of concentrated ultra-pure(up) nitric acid was added to 60µL samples. The same conditions for heat and ultrasonic bath as described above were applied. Then the samples were diluted to 300 µL by adding a solution of 1% (V/V) HNO_3_ and 30 µg/L Rh for an x5 total volume dilution. This gives a Rh concentration of 21 µg/L in the final sample.

Both sample series were analyzed using an Agilent Technologies 8800 ICP-QQQ-MS (Table S1 for instrumental parameters). The ICP-MS was fitted with a micro nebulizer with a flow of 50 µl/min to accommodate small sample volumes. The sample introduction was a high throughput setup. External standards containing Zn were used to calibrate the ICP-MS. Rh was added to the standards in the same concentration as the samples. Zn was analyzed in ammonia mode to remove any interferences on Zn, which was measured at 64 and 66 amu. Mass 64 has an isobaric overlap with ^64^Nickel (Ni); therefore, an inter-element correction for ^64^Ni interference on ^64^Zn was performed. The internal standard Rh was measured at 103 amu.

The method limit of detection and limit of quantification was calculated as 3 times the standard deviation of the buffer blank samples and 10 times the standard deviation of the buffer blank samples.

### Multiple sequence alignment

The sequences were aligned using Clustal Omega (McWilliam et al., 2013). The protein sequences used for the multiple sequence alignment were (UniProt ID): *Apis mellifera* (Q868N5), *Athalia rosae* (Q17083), *Pimpla nipponica* (O17428), *Pteromalus puparum* (B2BD67), *Encarsia formosa* (Q698K6), *Bombus ignites* (B9VUV6), *Bombus hypocrite* (C7F9J8), *Solenopsis invicta* (Q7Z1M0 and Q2VQM6), *Riptortus clavatus* (O02024), *Anthonomus grandis* (Q05808), *Lethocerus deyrollei* (B1B5Z4), *Aedes aegypti* (Q16927), *Nilaparvata lugens* (A7BK94), *Graptopsaltria nigrofuscata* (Q9U5F1), *Antheraea pernyi* (Q9GUX5), *Saturnia japonica* (Q59IU3), *Periplaneta Americana* (Q9U8M0), *Blattella germanica* (O76823), *Rhyparobia maderae* (Q5TLA5), *Homalodisca vitripennis* (Q0ZUC7), *Ichthyomyzon unicuspis* (Q91062), *Acipenser transmontanus* (Q90243), *Oreochromis aureus* (Q9YGK0), *Oncorhynchus mykiss* (Q92093), *Fundulus heteroclitus* (Q90508), *Xenopus laevis* (P18709), *Gallus gallus* (P87498), *Homo sapiens* (P55157) and *Anolis carolinensis* (Q9PUB1), based on Havukainen *et al*. (2011). The protein sequences used in the MSA with the CTCF protein were the same as listed above, but only including the β-barrel subdomain (residue 1 to 326 in honey bee Vg). The CTCF protein sequences used were: *Drosophila melanogaster* (Q9VS55), *Anopheles gambiae* (Q4G266), *Bombyx mori* (H9IXV8), *Tribolium castaneum* (D6WGY1), *Apis mellifera* (A0A7M7MWC7 and A0A7M7MWF4), and *Nasonia vitripennis* (A0A7M7H6S9). The CTCF proteins were identified through BLAST (Altschul et al., 1990), and the species represent the evolutionary relationship between *A. mellifera* and *D. melanogaster* (Honeybee Genome Sequencing, 2006). See Figure S5C for MSA.

### Identification of clusters

We used the full-length AlphaFold prediction of honey bee Vg for our structural analysis (Leipart et al., 2022). The first step was to identify conserved H and C residues (See Figure S1 for MSA). Second, we inspected their positions in 3D space to determine the distance and positions in relation to each other. Zinc usually coordinates with a minimum of four residues, so if our initial search did not identify at least four H/C in proximity, we went back to the MSA to identify adjacent (in sequence or 3D space) conserved S/D/E residues or regions of conserved residues (potential coordination through backbone association). Since the structural model lacks water molecules, we could not account for possible coordination involving hydrogen bonds from water molecules. The potential zinc-binding sites found in this way are called clusters. The MSA was uploaded to ConSurf (Ashkenazy et al., 2016) to create a PyMol script coloring atoms based on the degree of conservation. The results are presented in Figure S3. Finally, we evaluated the hydrophobic contrast of each cluster using Dudev and Lim’s (2003) formalism and using the PyMol command based on the Eisenberg hydrophobicity scale (Eisenberg et al., 1984). The results are presented in Figure S4. The *Apis mellifera* (Q868N5) sequence was input to ZincBind (Ireland and Martin, 2019), MotifScan (Pagni et al., 2007), and MOTIF Search (GenomeNet, Kyoto University Bioinformatics Center). Only MotifScan resulted in a positive hit.

### Limited proteolysis

A limited proteolysis analysis was performed to obtain the 40 kDa fragment of the β-barrel subdomain. Purified Vg was dephosphorylated with Lambda Protein Phosphatase (New England BioLabs, MA, USA). The samples were incubated with 1 µL Lambda Protein Phosphatase, 5 µL 10x NEBuffer for Protein MetalloPhosphatases and 5 µL of 10 mM MnCl_2_, for 30 minutes at 30°C. Then 6.5 ng of dephosphorylated Vg was digested with 5 and 10 units of caspase-1 (Sigma-Aldrich) for 2 hours at 37°C, and with 0.01, 0.1, and 1 unit of chymotrypsin (Sigma-Aldrich) for 30 minutes at 25°C. Two standards (ThermoFisher PageRuler™ unstained and pre-stained High Range Protein Ladder), unphosphorylated and undigested full-length Vg, and the digested samples were run on 4–20 % SDS-PAGE gel (Bio-Rad, CA, USA) under reducing conditions.

### Recombinant protein and preparation for element analysis

The β-barrel (amino acid 21 to 323) subdomain was produced by Genscript Biotech. The DNA was subcloned into a pET30a vector, with an N-terminal His tag and a SUMO solubility tag. The construct was expressed in E. coli Arctic Express (D3). A single colony was inoculated into an LB medium containing kanamycin, including 100 uM Zn^2+^. The bacteria were grown and harvested using standard approaches, and the target protein was resolubilized and purified using Ni-affinity purification. The tagged β-barrel subdomain was used in ICP-MS analysis. We made three sets of blank samples: buffer blank and two sets with SUMO-1 (human, His-tag, Enzo Life Sciences). One collection of SUMO-1 samples were incubated with 25uM ZnCl_2_ at 4 °C for 1 hour. The buffers in all four sample sets were exchanged to 0.5 M Tris HCl with 0.225 M NaCl. The ICP-MS protocol is explained above. ICP-MS revealed minimal Zn^2+^-coordination in all samples. The expression system was re-ordered; the LB medium included 42 uM Zn^2+^ or 50 uM Co^2+^, and one included both 42 uM Zn^2+^ and 50 uM Co^2+^. A SUMO-tag only expression system with the same three conditions as above was also ordered to avoid false positive results. We used the samples to perform UV-Vis spectroscopy, NMR spectroscopy and intrinsic tryptophan fluorescence spectroscopy for results and protocols (see supplementary.docx and Figure S8).

## Abbreviations

(DUF1943): Domain of unknown function
(ICP-MS): Inductively coupled plasma mass spectrometry
(MSA): multiple sequence alignment
(Vg): Vitellogenin
(vWF): von Willebrand factor

## Acknowledgment

The authors acknowledge The Research Council of Norway grant number 262137 for running costs and positions and BioCat (RCN grant number 249023) for travel grants and conference support.

## Data availability statement

All data presented here are included in the main article or the supplementary material (Supplementary.docx). The structural data for honey bee Vg was published earlier and supplemented there (Leipart et al., 2022).

## Author Contribution

VL executed structural analysis, purification, limited proteolysis and sample preparation for all experimental techniques. ØE performed ICP-MS analysis, VL and ØE analyzed the results. VL, DCT, OD, and ØH collaborated on the production of recombinant proteins. DCT and ØH performed UV-Vis spectroscopy, NMR spectroscopy, and intrinsic fluorescence spectroscopy. VL and FD analyzed motif data. ØH and GVA supervised the research. VL drafted the manuscript and made most figures. FD and ØH contributed with supplement figures. All authors contributed to writing the manuscript.

The authors declare no conflict of interest.

## References

Altschul, S. F., Gish, W., Miller, W., Myers, E. W. & Lipman, D. J. 1990. Basic local alignment search tool. J Mol Biol, 215, 403–10.

Amdam, G. V., Simoes, Z. L., Hagen, A., Norberg, K., Schroder, K., Mikkelsen, O., Kirkwood, T. B. & Omholt, S. W. 2004. Hormonal control of the yolk precursor vitellogenin regulates immune function and longevity in honeybees. Exp Gerontol, 39, 767–73.

Amdam, G. V., Aase, A. L., Seehuus, S. C., Kim Fondrk, M., Norberg, K. & Hartfelder, K. 2005. Social reversal of immunosenescence in honey bee workers. Exp Gerontol, 40, 939–47.

Anderson, T. A., Levitt, D. G. & Banaszak, L. J. 1998. The structural basis of lipid interactions in lipovitellin, a soluble lipoprotein. Structure, 6, 895–909.

Andreini, C., Banci, L., Bertini, I. & Rosato, A. 2006. Counting the Zinc-Proteins Encoded in the Human Genome. Journal of Proteome Research, 5, 196–201.

Ashkenazy, H., Abadi, S., Martz, E., Chay, O., Mayrose, I., Pupko, T. & Ben-Tal, N. 2016. ConSurf 2016: an improved methodology to estimate and visualize evolutionary conservation in macromolecules. Nucleic Acids Research, 44, W344–W350.

Ataie, N. J., Hoang, Q. Q., Zahniser, M. P. D., Tu, Y., Milne, A., Petsko, G. A. & Ringe, D. 2008. Zinc coordination geometry and ligand binding affinity: the structural and kinetic analysis of the second-shell serine 228 residue and the methionine 180 residue of the aminopeptidase from Vibrio proteolyticus. Biochemistry, 47, 7673–7683.

Auld, D. S., Falchuk, K. H., Zhang, K., Montorzi, M. & Vallee, B. L. 1996. X-ray absorption fine structure as a monitor of zinc coordination sites during oogenesis of Xenopus laevis. Proceedings of the National Academy of Sciences, 93, 3227–3231.

Babin, P. J., Bogerd, J., Kooiman, F. P., Van Marrewijk, W. J. A. & Van Der Horst, D. J. 1999. Apolipophorin II/I, Apolipoprotein B, Vitellogenin, and Microsomal Triglyceride Transfer Protein Genes Are Derived from a Common Ancestor. Journal of Molecular Evolution, 49, 150–160.

Baglivo, I., Russo, L., Esposito, S., Malgieri, G., Renda, M., Salluzzo, A., Di Blasio, B., Isernia, C., Fattorusso, R. & Pedone, P. V. 2009. The structural role of the zinc ion can be dispensable in prokaryotic zinc-finger domains. Proceedings of the National Academy of Sciences of the United States of America, 106, 6933–6938.

Baltaci, A. K. & Yuce, K. 2018. Zinc Transporter Proteins. Neurochemical Research, 43, 517–530.

Barozet, A., ChacÓN, P. & CortéS, J. 2021. Current approaches to flexible loop modeling. Current research in structural biology, 3, 187–191.

Biterova, E. I., Isupov, M. N., Keegan, R. M., Lebedev, A. A., Sohail, A. A., Liaqat, I., Alanen, H. I. & Ruddock, L. W. 2019. The crystal structure of human microsomal triglyceride transfer protein. Proceedings of the National Academy of Sciences, 116, 17251–17260.

Cassandri, M., Smirnov, A., Novelli, F., Pitolli, C., Agostini, M., Malewicz, M., Melino, G. & RaschellÀ, G. 2017. Zinc-finger proteins in health and disease. Cell Death Discovery, 3, 17071.

Chang, S., Jiao, X., Hu, J.-P., Chen, Y. & Tian, X.-H. 2010. Stability and folding behavior analysis of zinc-finger using simple models. International journal of molecular sciences, 11, 4014–4034.

Daniel, A. G. & Farrell, N. P. 2014. The dynamics of zinc sites in proteins: electronic basis for coordination sphere expansion at structural sites. Metallomics, 6, 2230–2241.

De Angelis, F., Lee, J. K., Connell, J. D., Miercke, L. J. W., Verschueren, K. H., Srinivasan, V., Bauvois, C., Govaerts, C., Robbins, R. A., Ruysschaert, J.-M., Stroud, R. M. & Vandenbussche, G. 2010. Metal-induced conformational changes in ZneB suggest an active role of membrane fusion proteins in efflux resistance systems. Proceedings of the National Academy of Sciences, 107, 11038.

Dudev, T. & Lim, C. 2003. Principles Governing Mg, Ca, and Zn Binding and Selectivity in Proteins. Chemical Reviews, 103, 773–788.

Eisenberg, D., Schwarz, E., Komaromy, M. & Wall, R. 1984. Analysis of membrane and surface protein sequences with the hydrophobic moment plot. J Mol Biol, 179, 125–42.

Falchuk, K. H. 1998. The molecular basis for the role of zinc in developmental biology. Mol Cell Biochem, 188, 41–8.

Fukada, T. & Kambe, T. 2011. Molecular and genetic features of zinc transporters in physiology and pathogenesis†. Metallomics, 3, 662–674.

Guidugli, K. R., Nascimento, A. M., Amdam, G. V., Barchuk, A. R., Omholt, S., Simoes, Z. L. & Hartfelder, K. 2005. Vitellogenin regulates hormonal dynamics in the worker caste of a eusocial insect. FEBS Lett, 579, 4961–5.

Gupta, G., Srivastava, P. P., Gangwar, M., Varghese, T., Chanu, T. I., Gupta, S., Ande, M. P., Krishna, G. & Jana, P. 2021. Extra-Fortification of Zinc Upsets Vitellogenin Gene Expression and Antioxidant Status in Female of Clarias magur brooders. Biological Trace Element Research.

Havukainen, H., Halskau, O., Skjaerven, L., Smedal, B. & Amdam, G. V. 2011. Deconstructing honeybee vitellogenin: novel 40 kDa fragment assigned to its N terminus. J Exp Biol, 214, 582–92.

Havukainen, H., Munch, D., Baumann, A., Zhong, S., Halskau, O., Krogsgaard, M. & Amdam, G. V. 2013. Vitellogenin recognizes cell damage through membrane binding and shields living cells from reactive oxygen species. J Biol Chem, 288, 28369–81.

Havukainen, H., Underhaug, J., Wolschin, F., Amdam, G. & Halskau, O. 2012. A vitellogenin polyserine cleavage site: highly disordered conformation protected from proteolysis by phosphorylation. J Exp Biol, 215, 1837–46.

Honeybee Genome Sequencing, C. 2006. Insights into social insects from the genome of the honeybee Apis mellifera. Nature, 443, 931–949.

Ilbert, M., Graf, P. C. & Jakob, U. 2006. Zinc center as redox switch--new function for an old motif. Antioxid Redox Signal, 8, 835–46.

Ireland, S. M. & Martin, A. C. R. 2019. ZincBind—the database of zinc binding sites. Database, 2019.

Jauch, R., Bourenkov, G. P., Chung, H.-R., Urlaub, H., Reidt, U., JäCkle, H. & Wahl, M. C. 2003. The Zinc Finger-Associated Domain of the Drosophila Transcription Factor Grauzone Is a Novel Zinc-Coordinating Protein-Protein Interaction Module. Structure, 11, 1393–1402.

Jernigan, R., Raghunathan, G. & Bahar, I. 1994. Characterization of interactions and metal ion binding sites in proteins. Current Opinion in Structural Biology, 4, 256–263.

Jumper, J., Evans, R., Pritzel, A., Green, T., Figurnov, M., Ronneberger, O., Tunyasuvunakool, K., Bates, R., ŽÍDek, A., Potapenko, A., Bridgland, A., Meyer, C., Kohl, S. A. A., Ballard, A. J., Cowie, A., Romera-Paredes, B., Nikolov, S., Jain, R., Adler, J., Back, T., Petersen, S., Reiman, D., Clancy, E., Zielinski, M., Steinegger, M., Pacholska, M., Berghammer, T., Bodenstein, S., Silver, D., Vinyals, O., Senior, A. W., Kavukcuoglu, K., Kohli, P. & Hassabis, D. 2021. Highly accurate protein structure prediction with AlphaFold. Nature, 596, 583–589.

Kluska, K., Adamczyk, J. & KrĘŻEl, A. 2018. Metal binding properties, stability and reactivity of zinc fingers. Coordination Chemistry Reviews, 367, 18–64.

Leipart, V., Montserrat-Canals, M., Cunha, E. S., Luecke, H., Herrero-Galan, E., Halskau, O. & Amdam, G. V. 2022. Structure prediction of honey bee vitellogenin: a multi-domain protein important for insect immunity. FEBS Open Bio, 12, 51–70.

Lim, M., Newman, J. A., Williams, H. L., Masino, L., Aitkenhead, H., Gravard, A. E., Gileadi, O. & Svejstrup, J. Q. 2019. A Ubiquitin-Binding Domain that Binds a Structural Fold Distinct from that of Ubiquitin. Structure, 27, 1316-1325.e6.

Liu, F., Su, Z., Chen, P., Tian, X., Wu, L., Tang, D.-J., Li, P., Deng, H., Ding, P., Fu, Q., Tang, J.-L. & Ming, Z. 2021. Structural basis for zinc-induced activation of a zinc uptake transcriptional regulator. Nucleic Acids Research, 49, 6511–6528.

Maksimenko, O. G., Fursenko, D. V., Belova, E. V. & Georgiev, P. G. 2021. CTCF As an Example of DNA-Binding Transcription Factors Containing Clusters of C2H2-Type Zinc Fingers. Acta Naturae, 13, 31–46.

Mariani, V., Biasini, M., Barbato, A. & Schwede, T. 2013. lDDT: a local superposition-free score for comparing protein structures and models using distance difference tests. Bioinformatics (Oxford, England), 29, 2722–2728.

Marreiro, D. D. N., Cruz, K. J. C., Morais, J. B. S., Beserra, J. B., Severo, J. S. & De Oliveira, A. R. S. 2017. Zinc and Oxidative Stress: Current Mechanisms. Antioxidants (Basel, Switzerland), 6, 24.

Martin, D. J. & Rainbow, P. S. 1998. The kinetics of zinc and cadmium in the haemolymph of the shore crab Carcinus maenas (L.). Aquatic Toxicology, 40, 203–231.

Matozzo, V., Gagné, F., Marin, M. G., Ricciardi, F. & Blaise, C. 2008. Vitellogenin as a biomarker of exposure to estrogenic compounds in aquatic invertebrates: A review. Environment International, 34, 531–545.

Mcwilliam, H., Li, W., Uludag, M., Squizzato, S., Park, Y. M., Buso, N., Cowley, A. P. & Lopez, R. 2013. Analysis Tool Web Services from the EMBL-EBI. Nucleic Acids Research, 41, W597–W600.

Mitchell, M. A. & Carlisle, A. J. 1991. Plasma zinc as an index of vitellogenin production and reproductive status in the domestic fowl. Comparative Biochemistry and Physiology Part A: Physiology, 100, 719–724.

Montorzi, M., Falchuk, K. H. & Vallee, B. L. 1994. Xenopus laevis vitellogenin is a zinc protein. Biochemical and biophysical research communications, 200, 1407–1413.

Pace, N. J. & Weerapana, E. 2014. Zinc-binding cysteines: diverse functions and structural motifs. Biomolecules, 4, 419–34.

Pagni, M., Ioannidis, V., Cerutti, L., Zahn-Zabal, M., Jongeneel, C. V., Hau, J., Martin, O., Kuznetsov, D. & Falquet, L. 2007. MyHits: improvements to an interactive resource for analyzing protein sequences. Nucleic Acids Res, 35, W433–7.

Pan, M. L., Bell, W. J. & Telfer, W. H. 1969. Vitellogenic Blood Protein Synthesis by Insect Fat Body. Science, 165, 393.

Perczel, A., GÁSpÁRi, Z. & Csizmadia, I. G. 2005. Structure and stability of β-pleated sheets*. Journal of Computational Chemistry, 26, 1155–1168.

Salmela, H., Harwood, G., MÜNch, D., Elsik, C., Herrero-GalÁN, E., Vartiainen, M. K. & Amdam, G. 2021. Nuclear Translocation of Vitellogenin in the Honey Bee (Apis mellifera). bioRxiv, 2021.08.18.456851.

Sappington, T. W. & S. Raikhel A., 1998. Molecular characteristics of insect vitellogenins and vitellogenin receptors. Insect Biochemistry and Molecular Biology, 28, 277–300.

Seehuus, S. C., Norberg, K., Gimsa, U., Krekling, T. & Amdam, G. V. 2006. Reproductive protein protects functionally sterile honey bee workers from oxidative stress. Proc Natl Acad Sci U S A, 103, 962–7.

Sevier, C. S. & Kaiser, C. A. 2002. Formation and transfer of disulphide bonds in living cells. Nature Reviews Molecular Cell Biology, 3, 836–847.

Sloup, V., JankovskÁ, I., NechybovÁ, S., PeŘInková, P. & LangrovÁ, I. 2017. Zinc in the Animal Organism: A Review. Scientia Agriculturae Bohemica, 48, 13–21.

Smolenaars, M. M. W., Madsen, O., Rodenburg, K. W. & Van Der Horst, D. J. 2007. Molecular diversity and evolution of the large lipid transfer protein superfamily^s⃞^. Journal of Lipid Research, 48, 489–502.

Stubbs L., Sun Y. & Caetano-Anolles D. 2011. Function and Evolution of C2H2 Zinc Finger Arrays.. In: T., H. (ed.) A Handbook of Transcription Factors. Subcellular Biochemistry. Springer, Dordrecht.

Sullivan, C. V. & Yilmaz, O. 2018. Vitellogenesis and Yolk Proteins, Fish. In: Skinner, M. K. (ed.) Encyclopedia of Reproduction (Second Edition). Oxford: Academic Press.

Tanaka, N., Kawachi, M., Fujiwara, T. & Maeshima, M. 2013. Zinc-binding and structural properties of the histidine-rich loop of Arabidopsis thaliana vacuolar membrane zinc transporter MTP1. FEBS Open Bio, 3, 218–224.

Tufail, M. & Takeda, M. 2008. Molecular characteristics of insect vitellogenins. Journal of Insect Physiology, 54, 1447–1458.

Zhang, T., Kuliyev, E., Sui, D. & Hu, J. 2019. The histidine-rich loop in the extracellular domain of ZIP4 binds zinc and plays a role in zinc transport. Biochem J, 476, 1791–1803.

Aase, A. L. T. O., Amdam, G. V., Hagen, A. & Omholt, S. W. 2005. A new method for rearing genetically manipulated honey bee workers. Apidologie, 36, 293–299.

