## Supplementary for "Where Honey Bee Vitellogenin may Bind Zn^2+^-Ions"

### Supplementary Material

#### Table S1. Instrumental parameters used for Agilent 8800 ICP-MS

| **ICP-MS** | **Settings[1]** | **Settings[2]** |
|---|---|---|
| RF power | 1600 W | 1600 W |
| Plasma gas | 15 L/min | 15 L/min |
| Auxiliary gas | 0,9 L/min | 0,9 L/min |
| Nebulizer gas | 0,90 L/min | 0,95 L/min |
| Makeup gas | 0,30 l/min | 0,20 l/min |
| Nebulizer pump | 0,32 rpm | 0,32 rpm |
| Sampler/skimmer | Ni | Ni |
| Sample Depth | 8,0 mm | 8,0 mm |
| **Data registration for ICP-MS** | | |
| Rinse time | 20 s | 20 s |
| Flush time | 50 s | 50 s |
| Read delay | 15 s | 15 s |
| Scanning mode | peak hop | peak hop |
| Points/spectral peak | 1 | 1 |
| Sweeps/reading | 10 | 10 |
| Replicates | 5 | 5 |
| P/A detector (puls/analog) | on | on |
| Temperature spray chamber | 12 °C | 2 °C |

1) First analysis series (full-length Vg)
2) Second analysis series (β-barrel subdomain)

### Supplementary methods for detection of zinc in the β-barrel subdomain

#### ICP-MS continued

We attempted to control for $Zn^{2+}$ interacting with the SUMO-tag by incubating the SUMO-tag with $Zn^{2+}$ before ICP-MS. Incubation resulted in significantly higher $Zn^{2+}$ levels in SUMO-tag samples compared to the non-incubated tagged β-barrel subdomain (Mann-Whitney U test: w = 0, p-value = 0.0114). The difference in concentration indicated that the incubation caused the association of $Zn^{2+}$ with the SUMO-tag. However, we could not rule out more specific $Zn^{2+}$-binding to SUMO. The net determined $Zn^{2+}$ concentration for the β-barrel subdomain was much lower compared to the concentration for native full-length Vg. Figure 1A shows a mean of 1524.0000 µg/L (SD ± 261.6562), while Figure S7 shows the mean as 64.2000 µg/L (SD ± 13.4350), and the significance suggested 0.01 bound $Zn^{2+}$ per β-barrel molecule. We attempted to examine this result with an independent approach and expressed the β-barrel subdomain in the presence of $Co^{2+,}$ attempting to displace $Zn^{2+}$ with this cation. $Co^{2+}$ is considered a good structural and functional model for studying $Zn^{2+}$-binding sites as the coordination is similar and exchange for $Zn^{2+}$ to $Co^{2+}$ occurs in nature (Lane and Morel, 2000, Shumilina et al., 2014). In contrast to $Zn^{2+}$, $Co^{2+}$ coordination causes a readily detected change in the protein's UV-Vis spectrum (Bertini and Luchinat, 1984, Shumilina et al., 2014, Sivo et al., 2017). The conducted experiments are presented below.

#### UV-Vis Spectroscopy

To identify the presence of $Zn^{2+}$-binding sites by $Co^{2+}$ substitution, SUMO fusion proteins were expressed and purified as described in the main manuscript (i.e., expressed in the presence of $Zn^{2+}$ [42 uM], $Zn^{2+}$ and $Co^{2+}$ [42 uM and 50 uM, respectively], and Co2+ [50 uM]).

Initially, in a storage buffer consisting of PBS at pH 7.4, 10% glycerol, and 0.5 M L-arginine, were centrifuged at 17000xg for 10 min. Supernatants were concentrated using Amicon filters with 10 kDa cut-off at 3250xg for 30 min, producing approximately 4 mg/mL protein samples. These were transferred to the UV-VIS cuvette (path length 10 mm, 500 uL capacity, Hellma item number 108-002-10-40). A UV-Vis spectra in the range of 200–800nm at room temperature was acquired. If $Co^{2+}$ successfully replaced $Zn^{2+}$, we would have expected to see a distinct $Co^{2+}$-specific peak pattern near the 500–750 nm range (Sivo et al., 2017, Lane and Morel, 2000). However, we did not observe this (negative results are presented in Figure S8A). The samples were then used for NMR and intrinsic tryptophan fluorescence spectroscopy (see below).

### NMR Spectroscopy

$Co^{2+}$ can create a paramagnetic shift in protein NMR spectra (Lane and Morel, 2000). Therefore, we transferred stocks of SUMO-fusion proteins exposed to $Zn^{2+}$, $Zn^{2+}$ $Co^{2+}$, and $Co^{2+}$ (as described above) to NMR tubes. $D_2O$ up to 5% v/v was added. We then acquired 1D proton spectra (25°C, water suppression using Watergate, 512 scans, processed using exponential multiplication with a line broadening of 0.3 Hz), and inspected the spectra without finding any significant differences between the samples (Figure S8B).

### Intrinsic Tryptophan Fluorescence Spectroscopy

We also looked for fold changes in response to divalent cation(s) present during expression. To do this, we transferred 5 uL of the NMR samples (described above) into 300 PBS buffer at pH 6.5 in a quartz cuvette (path length 5 mm) and conducted an emission scan (excitation wavelength 295 nm, slit widths 5 nm, 310–500 nm). The spectra, which primarily provide information on the microenvironment of the tryptophans in the protein (Knappskog and Haavik, 1995, Takita et al., 2003) did not show any significant difference.

### Figure legends

**Figure S1: Multiple sequence alignment.** Snapshots of the multiple sequence alignment. All the included species are noted with their UniProt ID. Residues included in clusters are in bold colors (αh.1: green, αh.2: red, Duf.1: dark blue, Duf.2: orange, Ct: yellow, βb.1: pink and βb.2: cyan). Some alignment regions are excluded (noted with "…") since they are not relevant to this study or have significant gaps. **A)** The residues from βb.1, βb.2, and one residue from cluster αh.1 are in the β-barrel subdomain. In addition, the DNA binding motif (pink box) is part of this subdomain. The conserved residues are colored (bold black). **B)** Cluster αh.1, αh.2, Duf.1 and Duf.2 are in the lipid binding site (DUF1943). In addition, the suggested zinc coordinating residues from studies in Lamprey (gray bold) and the MotifScan zinc-binding site (yellow box) are found in this region. H926 is colored (black bold). **C)** No cluster was identified in the region before the vWF domain, but the conserved disulfide bridges residues are colored (brown bold). **D)** Cluster Ct is in the C-terminal region, where all the conserved C and H residues are marked.

**Figure S2: MotifScan.** The zinc-binding motif (yellow) identified by MotifScan is located in the DUF1943 domain (purple) but extends into the cavity of the β-barrel (green). The predicted zinc-binding H residue (H926) is shown as a stick. Cluster Duf.1 (blue spheres), βb.1 (magenta spheres), βb.2 (cyan spheres), H229 from αh.1 (green stick), and the DNA binding motif (pink) is in close proximation to this predicted zinc-binding motif.

**Figure S3: Conservation.** Residues are colored using ConSurf, based on the MSA. The low to highly conserved residues are colored from light blue to dark pink (scale presented in the lower-right corner). Both the secondary structures and the spheres (representing the clusters) are colored according to this scale. **A)** The buried residues in the β-barrel subdomain are well conserved, including the residues in cluster βb.1, βb.2, and the DNA binding motif β-sheet. The regions closer to the surface are less conserved. **B)** The α-helices in the α-helical subdomain are well conserved. The residues in Cluster αh.1 and αh.2 are also conserved. **C)** One of the β-sheet in the DUF1943 domain includes cluster Duf.1, Duf.2, and the zinc motif identified by MotifScan. The conservation of the residues in the β-sheet are variable, but the clusters and zinc motif are conserved. As shown in the MSA, residue H926 is not conserved. **D)** The C-terminal is generally not conserved, except the four residues presented as cluster Ct and the third disulfide bridge (labeled).

**Figure S4: Hydrophobicity.** The surface and secondary structure are colored using the Eisenberg hydrophobicity scale (scale presented in the lower-left corner). In both panels, the clusters are shown as spheres and marked with a blue dotted circle. Their respective domains are presented as surface and cartoon. **A)** Cluster αh.2 is the position between the highly hydrophobic core of the lipid binding site (β-sheet) and the polar surface of the α-helical subdomain. **B)** Cluster Duf.1 and Duf.2 are positioned on the same β-sheet that make up one side of the lipid binding site (highly hydrophobic), while the other side is facing the surface and is more polar.

**Figure S5: Logo representation of DNA binding site motifs and sequence analysis of CTCF. A)** The most significant motif (motif A) found by Salmela *et al.* (2021) for Vg-DNA binding sites, as shown in Figure 5. **B)** The motif for CTCF in Drosophila melanogaster from the JASPAR database (Castro-Mondragon et al., 2021) (matrix MA0531.1). The logo representation was made with WebLogo3 (Schneider and Stephens, 1990, Crooks et al., 2004). **C)** Residue 140 to 233 in the β-barrel subdomain, using the same species as in the full-length MSA (Figure S1), aligned to CTCF proteins. The conserved residues from the β-barrel subdomain, identified in the CTCF proteins, are in bold.

**Figure S6. Proteolysis of honey bee Vg by caspase-1 and chymotrypsin.** The black arrow emphasizes the probable 40 kDa cleavage products of the full-length Vg (flVg) with 5 and 10 units of caspase-1 in lanes 3 and 4, respectively. We did not identify a clear 150 kDa band. The smaller bands outside the range of the standard could be the lambda protein phosphates (25 kDa) or caspase-1 (30 kDa). The chymotrypsin (lane 5 to 7) cleaves Vg completely into small fragments, and no 40 kDa band was identified. The smaller bands outside the range of the standard could be lambda protein phosphates (25 kDa) or chymotrypsin (25 kDa).

**Figure S7. ICP-MS results for the β-barrel subdomain.** The concentration was measured with ICP-MS for the x5 samples of SUMO tagged β-barrel subdomain (bb), sample buffer (blk), non-incubated SUMO tag (Sblk), and SUMO-tag incubated with $Zn^{2+}$ (SZnblk). The mean and the standard deviation of the mean are indicated for each group.

**Figure S8. Spectroscopic analyses of SUMO-fusion proteins expressed in $Zn^{2+}$, $Zn^{2+}$ and $Co^{2+}$, and $Co^{2+}$ medium. A)** UV-Vis spectra of protein expressed in medium enriched with $Zn^{2+}$ (green traces), $Zn^{2+}$ and $Co^{2+}$(blue traces), and $Co^{2+}$ (red traces). Expressions of the Sumo tag only are represented by dashed lines, and the SUMO beta-barrel fusion protein is represented by whole lines. **B)** Amide region from $^{1}H$ NMR spectra of SUMO beta-barrel fusion protein expressed in medium enriched with $Zn^{2+}$ (green trace), $Zn^{2+}$ and $Co^{2+}$(blue trace), and $Co^{2+}$ (red trace). **C)** Intrinsic tryptophan fluorescence spectra of SUMO beta-barrel fusion protein expressed in medium enriched with $Zn^{2+}$ (green trace), $Zn^{2+}$ and $Co^{2+}$(blue traces), and $Co^{2+}$ (red trace).

### References


BERTINI, I. & LUCHINAT, C. 1984. High spin cobalt(II) as a probe for the investigation of metalloproteins. *Adv Inorg Biochem,* 6**,** 71-111.

CASTRO-MONDRAGON, J. A., RIUDAVETS-PUIG, R., RAULUSEVICIUTE, I., BERHANU LEMMA, R., TURCHI, L., BLANC-MATHIEU, R., LUCAS, J., BODDIE, P., KHAN, A., MANOSALVA PÉREZ, N., FORNES, O., LEUNG, TIFFANY Y., AGUIRRE, A., HAMMAL, F., SCHMELTER, D., BARANASIC, D., BALLESTER, B., SANDELIN, A., LENHARD, B., VANDEPOELE, K., WASSERMAN, W. W., PARCY, F. & MATHELIER, A. 2021. JASPAR 2022: the 9th release of the open-access database of transcription factor binding profiles. *Nucleic Acids Research*.

CROOKS, G. E., HON, G., CHANDONIA, J. M. & BRENNER, S. E. 2004. WebLogo: a sequence logo generator. *Genome Res,* 14**,** 1188-90.

KNAPPSKOG, P. M. & HAAVIK, J. 1995. Tryptophan Fluorescence of Human Phenylalanine Hydroxylase Produced in Escherichia coli. *Biochemistry,* 34**,** 11790-11799.

LANE, T. W. & MOREL, F. M. 2000. Regulation of carbonic anhydrase expression by zinc, cobalt, and carbon dioxide in the marine diatom Thalassiosira weissflogii. *Plant physiology,* 123**,** 345-352.

SCHNEIDER, T. D. & STEPHENS, R. M. 1990. Sequence logos: a new way to display consensus sequences. *Nucleic Acids Res,* 18**,** 6097-100.

SHUMILINA, E., DOBROVOLSKA, O., DEL CONTE, R., HOLEN, H. W. & DIKIY, A. 2014. Competitive cobalt for zinc substitution in mammalian methionine sulfoxide reductase B1 overexpressed in E. coli: structural and functional insight. *Journal of biological inorganic chemistry : JBIC : a publication of the Society of Biological Inorganic Chemistry,* 19**,** 85-95.

SIVO, V., D'ABROSCA, G., RUSSO, L., IACOVINO, R., PEDONE, P. V., FATTORUSSO, R., ISERNIA, C. & MALGIERI, G. 2017. Co(II) Coordination in Prokaryotic Zinc Finger Domains as Revealed by UV-Vis Spectroscopy. *Bioinorg Chem Appl,* 2017**,** 1527247.

TAKITA, T., NAKAGOSHI, M., INOUYE, K. & TONOMURA, B. I. 2003. Lysyl-tRNA Synthetase from Bacillus stearothermophilus: The Trp314 Residue is Shielded in a Non-polar Environment and is Responsible for the Fluorescence Changes Observed in the Amino Acid Activation Reaction. *Journal of Molecular Biology,* 325**,** 677-695.

Figure S1

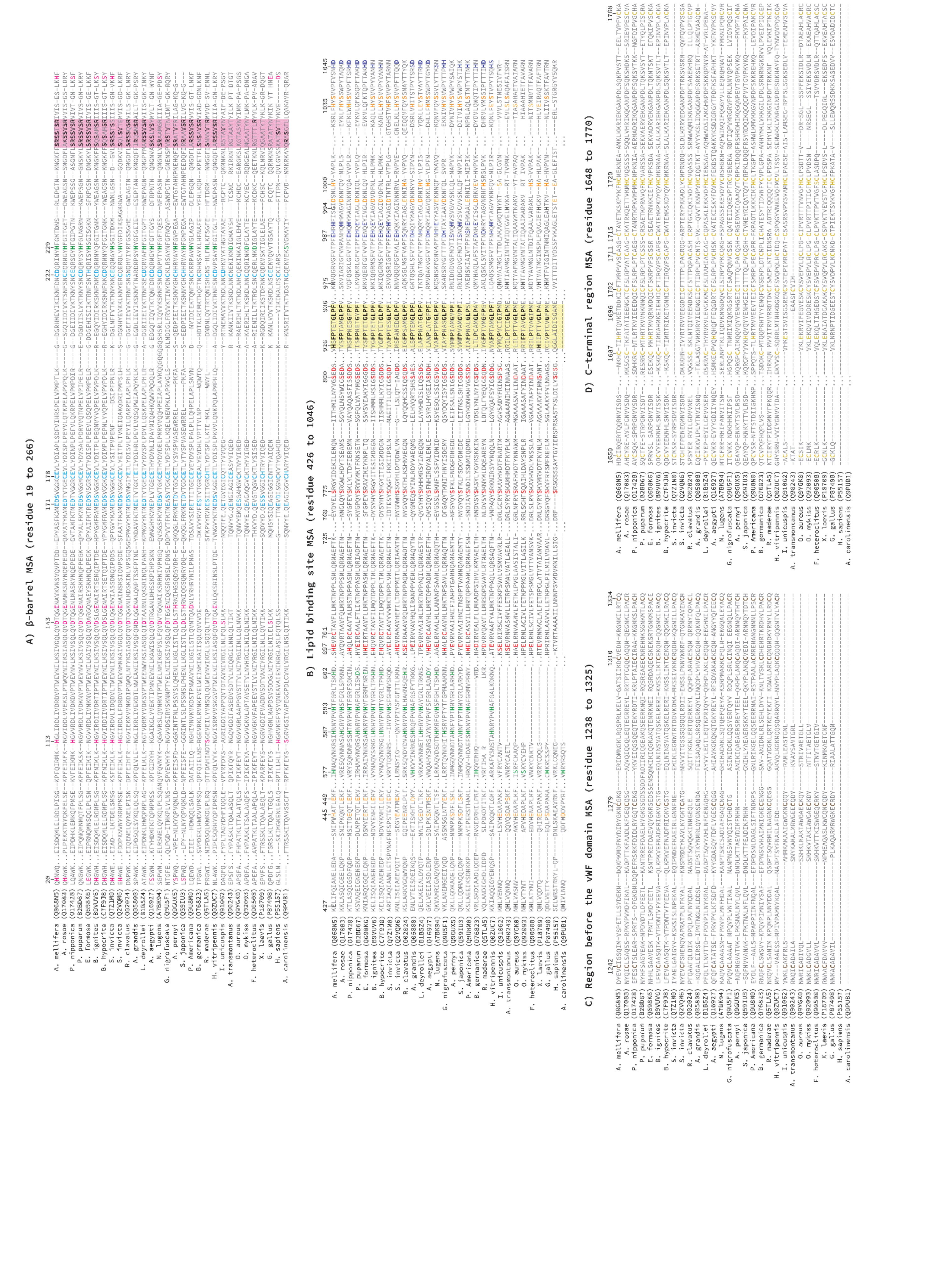

A) β-barrel MSA (residue 19 to 266)
B) Lipid binding site MSA (residue 426 to 1046)
C) Region before the vWF domain MSA (residue 1238 to 1325)
D) C-terminal region MSA (residue 1648 to 1770)

Figure S2

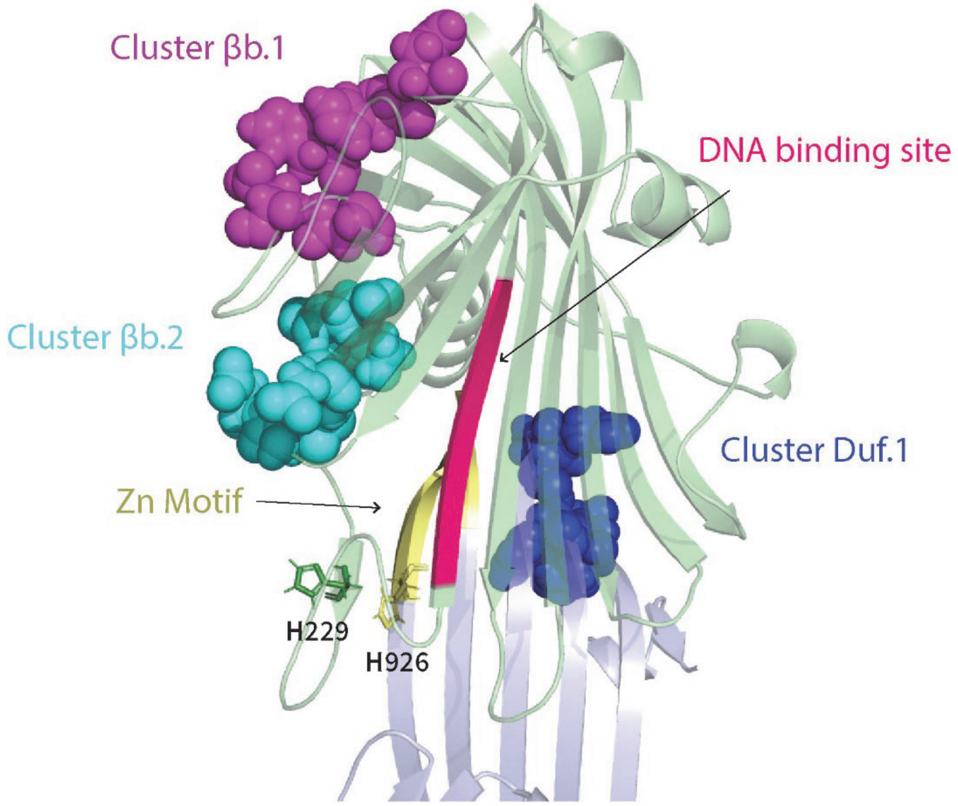

Cluster βb.1
DNA binding site
Cluster βb.2
Cluster Duf.1
Zn Motif
H229
H926

Figure S3

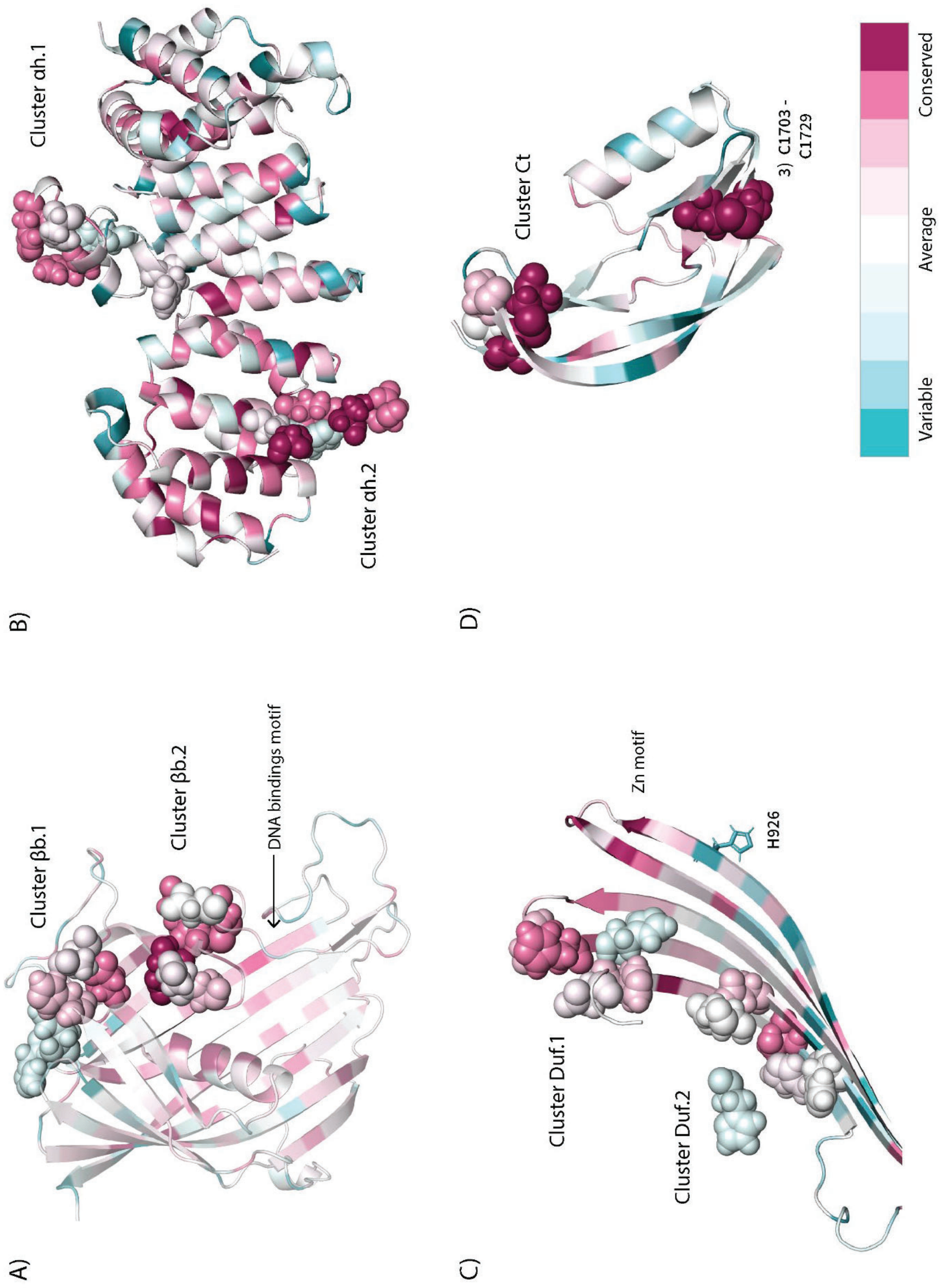
A)
Cluster βb.1
Cluster βb.2
DNA bindings motif
B)
Cluster αh.1
Cluster αh.2
C)
Cluster Duf.1
Cluster Duf.2
Zn motif
H926
D)
Cluster Ct
3) C1703 -
C1729
Variable
Average
Conserved

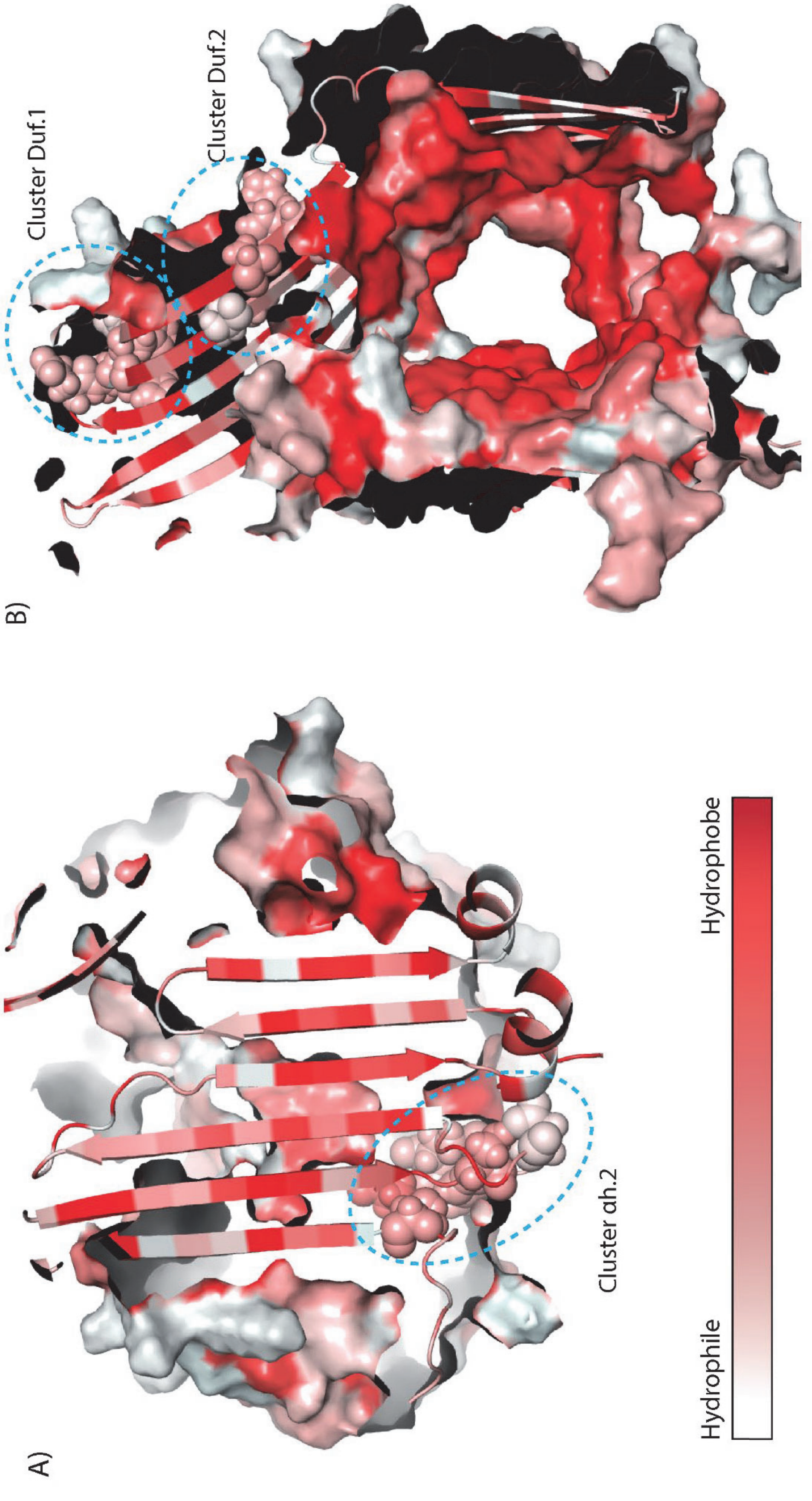
A)
B)
Cluster Duf.1
Cluster Duf.2
Cluster ah.2
Hydrophile
Hydrophobe

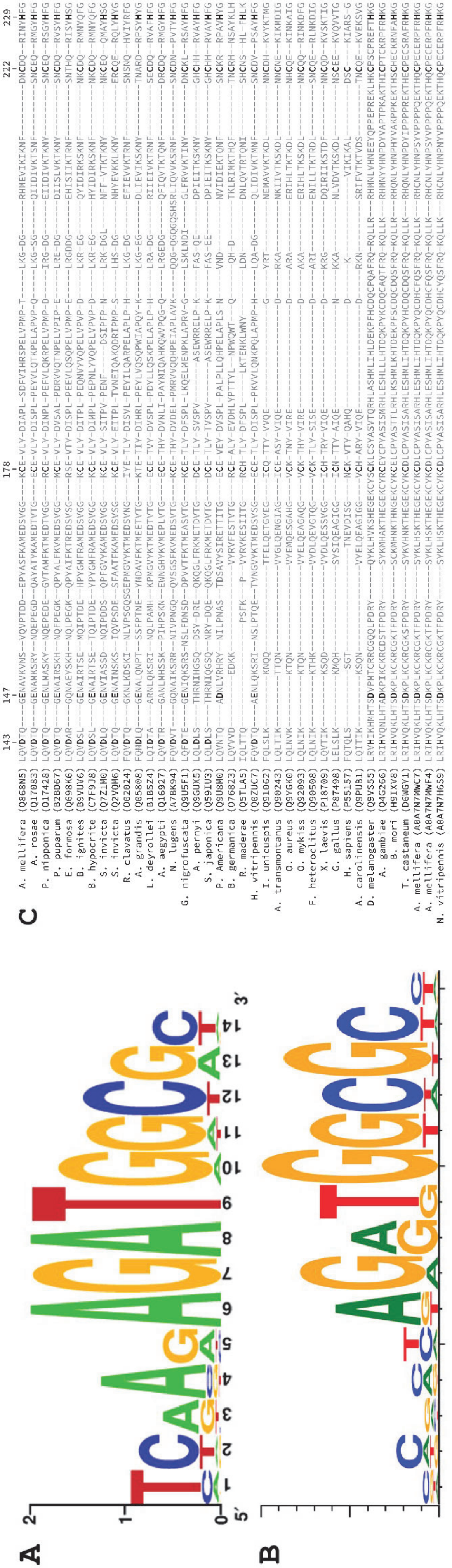
A
B
C

Figure S6

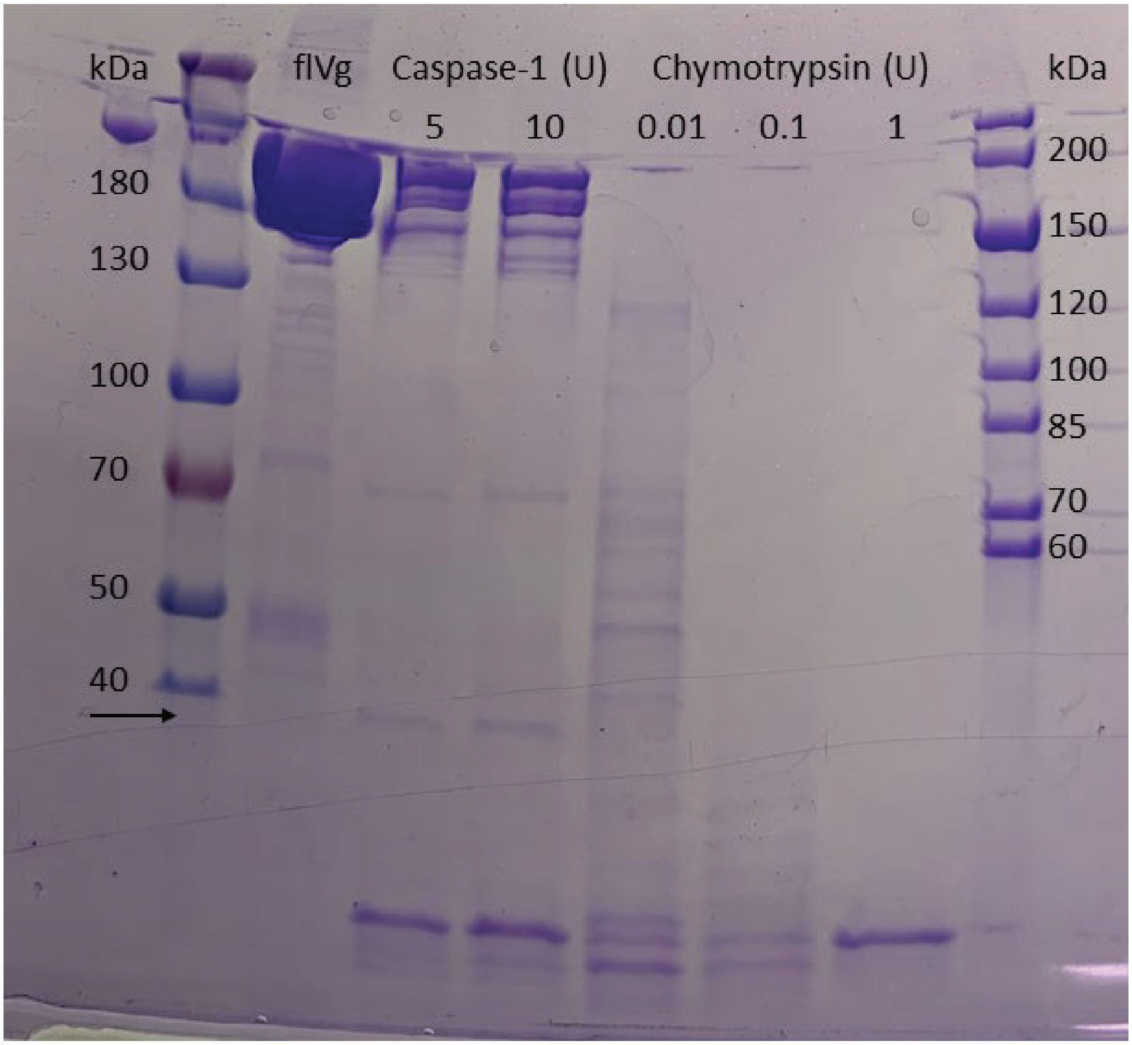
kDa
flVg
Caspase-1 (U)
Chymotrypsin (U)
kDa
5
10
0.01
0.1
1
180
130
100
70
50
40
200
150
120
100
85
70
60

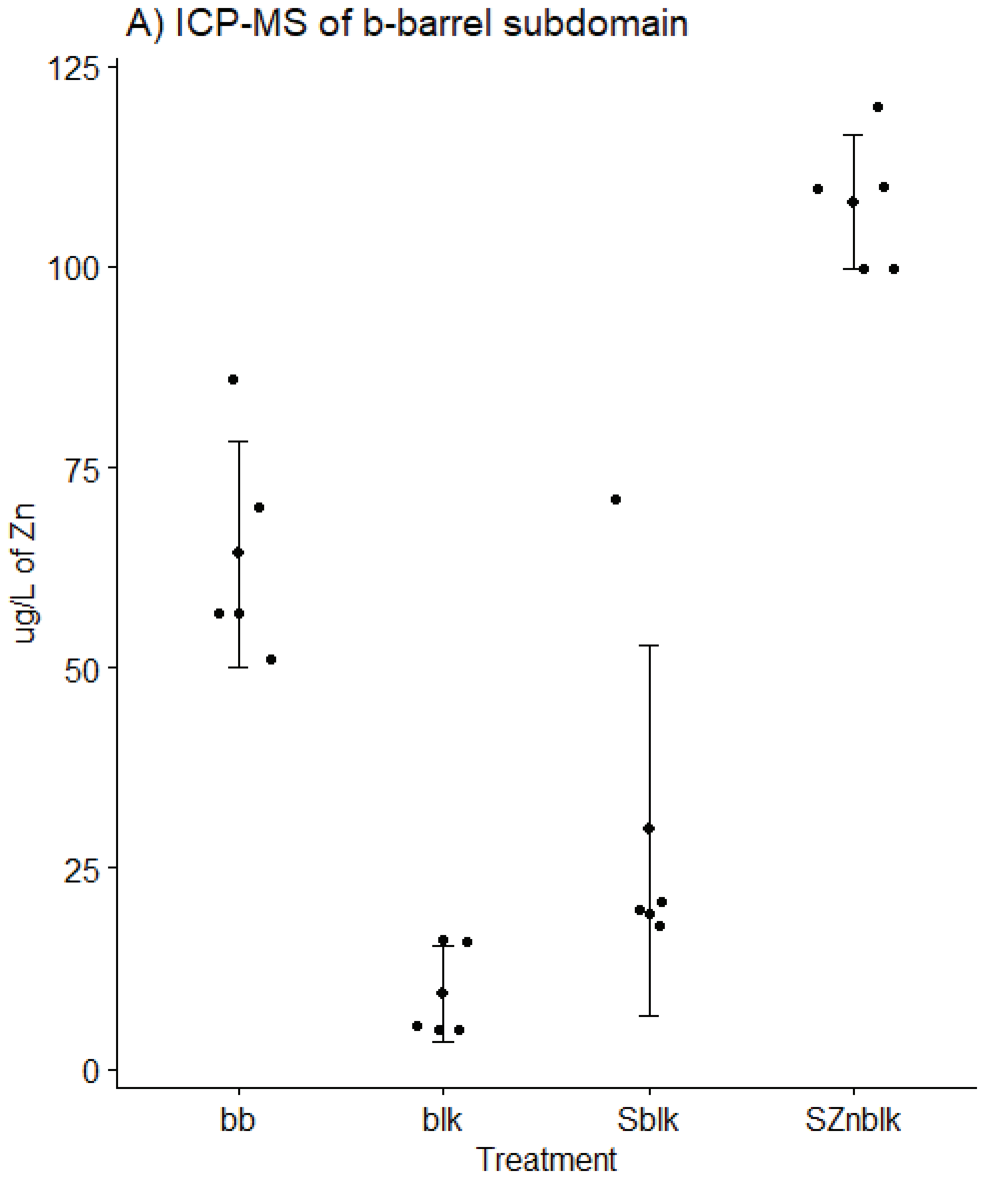
A) ICP-MS of b-barrel subdomain
125
100
75
50
25
0
ug/L of Zn
bb
blk
Sblk
SZnblk
Treatment

Figure S8

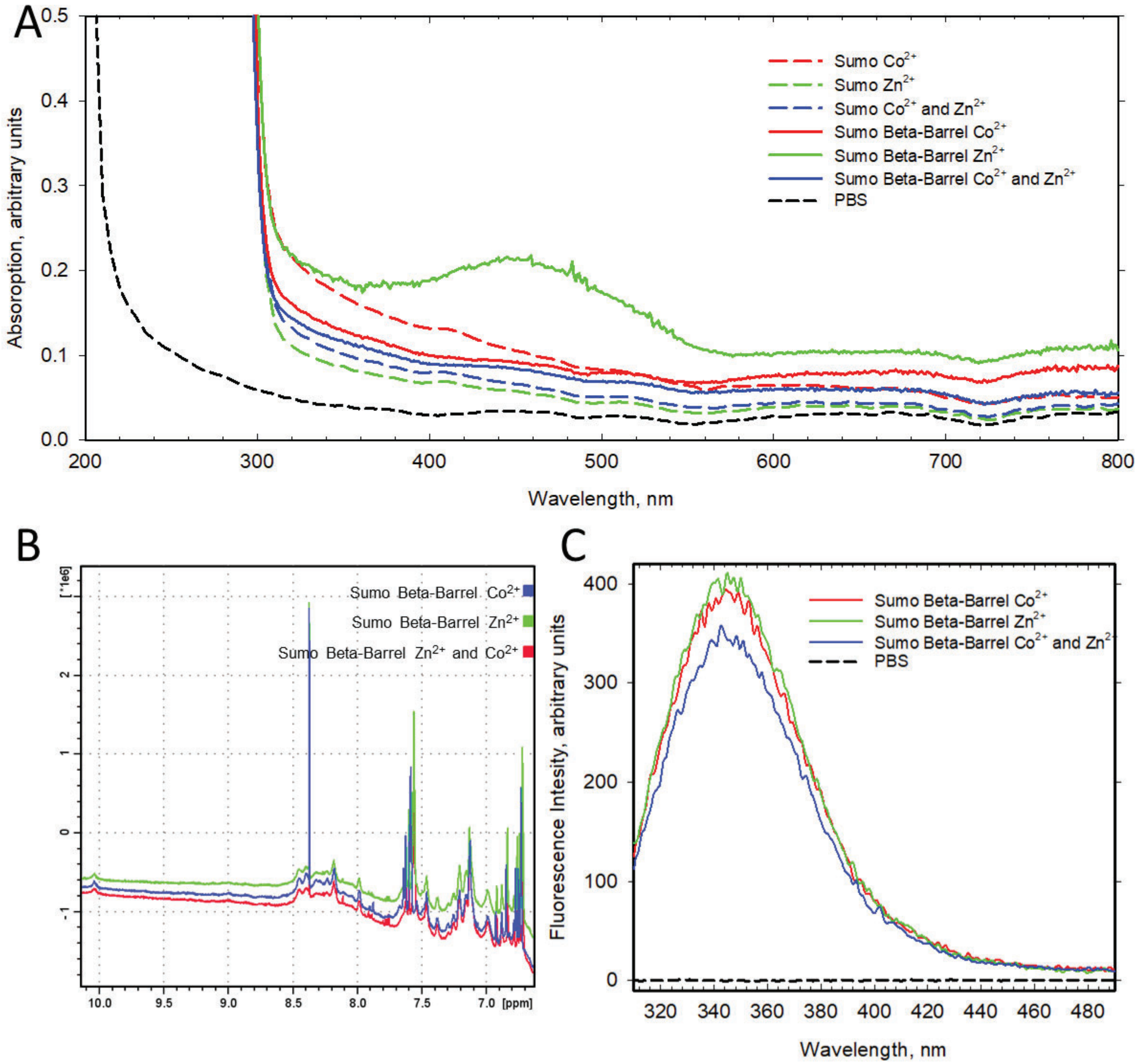
A
Absoroption, arbitrary units
Wavelength, nm
Sumo Co2+
Sumo Zn2+
Sumo Co2+ and Zn2+
Sumo Beta-Barrel Co2+
Sumo Beta-Barrel Zn2+
Sumo Beta-Barrel Co2+ and Zn2+
PBS
B
[*1e6]
[ppm]
Sumo Beta-Barrel Co2+
Sumo Beta-Barrel Zn2+
Sumo Beta-Barrel Zn2+ and Co2+
C
Fluorescence Intesity, arbitrary units
Wavelength, nm
Sumo Beta-Barrel Co2+
Sumo Beta-Barrel Zn2+
Sumo Beta-Barrel Co2+ and Zn2+
PBS
